# Preservation of an Aging-Associated Mitochondrial Signature in Advanced Human Neuronal Models

**DOI:** 10.1101/2024.03.28.587193

**Authors:** Nimmy Varghese, Leonora Szabo, Zameel Cader, Imane Lejri, Amandine Grimm, Anne Eckert

**Affiliations:** Research Cluster Molecular and Cognitive Neurosciences, University of Basel, 4002 Basel, Switzerland; Neurobiology Lab for Brain Aging and Mental Health, University Psychiatric Clinics Basel, 4002 Basel, Switzerland; Nuffield Department of Clinical Neuroscience, University of Oxford, Oxford, UK

**Keywords:** Aging, induced pluripotent stem cells-derived neurons, directly converted neurons, mitochondria, glycolysis

## Abstract

This study investigated whether induced pluripotent stem cell-derived neurons (iPSCsNs) and directly converted neurons (iNs) generated from the same cells of origin (human fibroblasts) represent aging-related characteristics on mitochondrial levels. There is still uncertainty regarding the potential for rejuvenation or preservation of an aging-associated donor signature in aged iPSCsNs upon transition through pluripotent states, while direct conversion retains the aging-associated mitochondrial impairments. Surprisingly, both aged neuronal models exhibited age-associated donor phenotypes, including decreased ATP, mitochondrial membrane potential, mitochondrial respiration, NAD^+^/NADH ratio, and increased radical levels and mitochondrial mass. Besides, a fragmented mitochondrial network was observed in both aged neuronal models. However, unlike aged iNs, aged iPSCsNs did not show a metabolic shift towards anaerobic glycolysis to compensate for the energy deficit. Moreover, the mRNA expression profile significantly differed between aged iPSCsNs and aged iNs. Our study indicates that aged iPSCsNs may experience rejuvenation in certain parameters, such as transcriptomics and the aging-associated glycolytic shift. Nevertheless, aged iPSCsNs can be a valuable tool for studying neuronal aging of mitochondrial parameters *in vitro* alongside aged iNs.

## 3. Introduction

Life expectancy has dramatically increased in recent decades. Nowadays, everyone is expected to live up to 60 years or more [1, 2]. However, the increase in life expectancy is accompanied with age-related challenges, including cognitive deterioration and an increased susceptibility to neurodegenerative conditions [3]. In order to tackle this issue, the World Health Organization has officially designated the period spanning from 2021 to 2030 as the "United Nations Decade of Healthy Aging". The primary objective of this initiative is to enhance the overall well-being of aged individuals [4]. In the interest of promoting healthy brain aging and enhancing our knowledge of neurodegenerative diseases, it is crucial to investigate the predisposing molecular factors of brain aging.

The understanding of aging, particularly in the human brain at the molecular and cellular level, is still poorly investigated. A comprehensive understanding of the biological processes behind aging is crucial in achieving the goal of optimal brain health. Aging is a complex phenomenon marked by a gradual decline in physiological function and behavioral capacity. This natural process emerges from the gradual accumulation of molecular and cellular damage during an individual’s lifetime [5, 6]. The accumulation of damage might ultimately surpass the body’s capacity to repair and eliminate impaired materials, resulting in an imbalance that hampers the optimal functioning of different body systems. This time-dependent accumulation of impairments can increase the vulnerability to develop neurodegenerative diseases. The molecular and cellular aging process is a complex field with many unsolved questions regarding the underlying mechanisms of aging-associated alterations. As a result, researchers are directing their efforts toward comprehending the fundamental mechanism of aging to counteract aging manifestation and prevent the onset of neurodegenerative disorders [7]. Understanding the alteration on the molecular and cellular level of aging is diverse and challenging to characterize. In the pioneering overview of Lopez et al. [8], twelve hallmarks were identified to describe the cellular and molecular alteration of aging, including mitochondrial dysfunction and telomere attrition. As research on aging continues to evolve, we can expect to discover additional hallmarks contributing to the aging process. In aging, energy impairments and imbalances between the demand and supply of energy are often closely associated [9]. Mitochondria are essential for sustaining life by facilitating the energy conversion processes within the cells. Consequently, impaired mitochondrial functioning can severely disrupt the cellular energy balance, with neurons being the most vulnerable cells to aging due to their heavy reliance on the energy generated by mitochondria [10–13]. Gaining insight into the mechanisms behind mitochondrial malfunction in the context of aging and age-related metabolic disorders has significant potential for advancing our understanding and facilitating the development of therapeutic interventions aimed at mitigating the adverse health outcomes associated with aging.

Still, studying aging, particularly brain aging, poses numerous unanswered questions regarding its underlying mechanism. To gain a better knowledge of age-related changes and develop new therapeutic strategies for prevention, it is essential to find and utilize suitable *in vitro* models that accurately mimic *in vivo* human brain aging. Cultures of neuronal cells are crucial to understanding the nervous system. However, practical tools to investigate human neuronal aging *in vitro* are limited [14, 15]. The inaccessibility and ethical constraints of viable human neuronal cells further restrict the investigation of the precise mechanism underlying the aging brain. To this main struggle, the research relies heavily on animal models, which are still an essential tool in the aspect of gene-to-behavior interaction. Nevertheless, animal models present a major obstacle: animal models are not human, especially in the context of transferability to the human brain’s architecture, development, and cognitive ability [16]. The effectiveness of treatment strategies predicted by animal experiments remains controversial due to their failure to translate to clinical investigation, especially on disease-related levels [17, 18]. In general, the research and development of pharmaceutical interventions targeting aging and neurodegenerative illnesses pose significant challenges due to the limited availability of healthy, viable human neuronal cells [19]. In recent years, new approaches have emerged for generating neuronal cells, such as through nuclear reprogramming of somatic cells [20, 21]. This includes the advanced neuronal models of induced pluripotent stem cell-derived neurons (iPSCsNs) [22] and direct conversion of neurons (iNs) from human fibroblasts (HFs) [23, 24]. These advanced reprogramming technologies opened the possibility of obtaining human neurons from more easily accessible somatic cell lines, providing scientists with effective models to understand the molecular processes underlying neuronal functions and develop new pharmaceutical interventions. Although more protocols are available for generating iPSCsNs than directly converted neurons, both model systems have advantages and limitations. As a scientist, research budgets are often limited, making time and money crucial when choosing *in vitro* models for investigation [25, 26]. Moreover, a model system that yields a high number of cells is preferable. When comparing the generation of iNs and iPSCsNs, it is evident that iNs require less time to generate by skipping the pluripotent state [21]. On the other hand, iPSCsNs need a longer generation time as they undergo an embryonic-like state. Once established, iPSCs can be expanded infinitely, providing an abundant supply of stable cells. In many protocols, iPSCs differentiation involves generating a stable and expandable neuronal progenitor stage that serves as an intermediate cell line before differentiation into neurons. Producing a substantial quantity of neurons via iPSCs differentiation is highly advantageous for conducting bioenergetic studies, drug screening, and transplantation, as these applications require a considerable amount of material. In comparison, generating iNs takes fewer weeks but is limited by the number of HFs used, making extensive screenings challenging. Nevertheless, the main factor in the interest in choosing the cellular models is how well the model systems, after reprogramming, still represent the aging- or disease-associated phenotype. From that point of view, the use of iPSCs technology seemed to be constrained by the process of cellular rejuvenation leading to a reset of age indicators from the original donor cells [20, 25, 27–29]. The process of direct neural reprogramming entails the conversion of a somatic cell into a neuron without the need for an intermediary pluripotent stage [28, 30–33]. By skipping the proliferative or stem-cell-like stage (rejuvenation), the iNs are considered to preserve their age-associated hallmarks from the initial somatic cell source [23, 24, 34–36]. Hereby, the direct conversion of fibroblasts into induced neurons (iNs) represents an alternative avenue for generating human neurons *in vitro* by preserving an aging phenotype [27, 37, 38]. Thus, preservation of human age seems unlikely in the iPSCsNs, given that these cells must transit the embryo-like iPSCs state, which likely rejuvenates somatic cells of aged donors [37, 39]. Still, in recent years, some papers have arisen tackling the question of how well the iPSCs and their derived cell are really losing the aging donor signature [40, 41].

In our previous study (unpublished data of the Eckert group), we demonstrated that aging-associated phenotypes on the mitochondrial level could be observed in aged iPSCs by comparing them to young iPSCs. This raises the question of whether the pluripotent state of iPSCs is undergoing rejuvenation. In this mind, it is possible that aged iPSCsNs could also represent an aging phenotype or that the iNs model system is the only viable option for the aging research. The primary objective of this study was to examine the extent to which the advanced neuronal models of iPSCsNs and iNs derived from aged human donors accurately represent the aging-related characteristics of their donor cells by analyzing alterations on the mitochondrial level. Our research revealed that iPSCsNs derived from aged individuals displayed a comparable mitochondrial phenotype to that of aged iNs from the same donors. This paper highlights an aging-associated donor phenotype at the mitochondrial level in aged iPSCsNs, which contradicts an overall rejuvenation mechanism. The findings open new possibilities in a field with intense controversy. Overall, this study deepened our understanding of how well iNs or iPSCsNs from aged donors mirror features of human brain aging by modeling the aging-related changes *in vitro*.

## 4. Results

### 4.1. Assessing the mitochondrial impairment in aged HFs compared to young HFs

As part of our investigation, we first examined the aging signatures of the corresponding donor HFs, which were then further converted to iPSCsNs or iNs. By comparing HFs from four young (Age_mean_ = 31 years, SD = 5.03) and four aged (Age_mean_ = 69 years, SD = 7.632) individuals, we identified an aging signature related to mitochondrial properties. These defined properties in aged HFs were used to determine the aging-associated phenotype, which we then attempted to observe in the iNs and iPSCsNs generated from the HFs of the same donors.

A deficit in the bioenergetic state of aging is mainly linked to mitochondrial impairments [42]. First, we measured the cellular adenosine triphosphate (ATP) level (Fig 1A) in the HFs to determine the metabolic activity in the whole cell. We found a significant decline in the overall ATP level in the aged HFs compared to the young HFs. Following, we detected the mitochondrial membrane potential (MMP or ΔΨ_m_), which is the driving force of Complex V, the ATPase, in the ETC [43]. The MMP (Fig 1B) in aged HFs showed a drop in comparison to the young HFs. Observing decreased MMP in age can be another indicator of impaired mitochondrial health. To receive a more precise image of the mitochondrial respiratory activity, mainly associated with oxidative phosphorylation (OXPHOS), the oxygen consumption rate (OCR) was measured using the Seahorse analyzer to perform the Seahorse XF Cell Mito Stress Test (Fig 1C & E). The aged HFs represented a lower mitochondrial respiratory rate (Fig 1C), indicated by a lower OCR. Moreover, in all determined mitochondrial bioenergetic parameters (Fig 1E) calculated from the OCR profile, basal respiration, ATP-production coupled respiration, maximal respiration, and spare respiration capacity indicated a significant loss of respiratory properties. Besides mitochondrial respiration, glycolysis is the second primary energy source in the cells [44], which was assessed using the Seahorse XF Glycolysis Stress Test Kit (Fig 1D & E). By sequential injection of different molecules, the glycolysis parameters glycolysis and glycolysis capacity were determined by measuring the extracellular acidification rate (ECAR). We thereby demonstrated a substantial increase in all glycolytic parameters.

**Figure 1:**
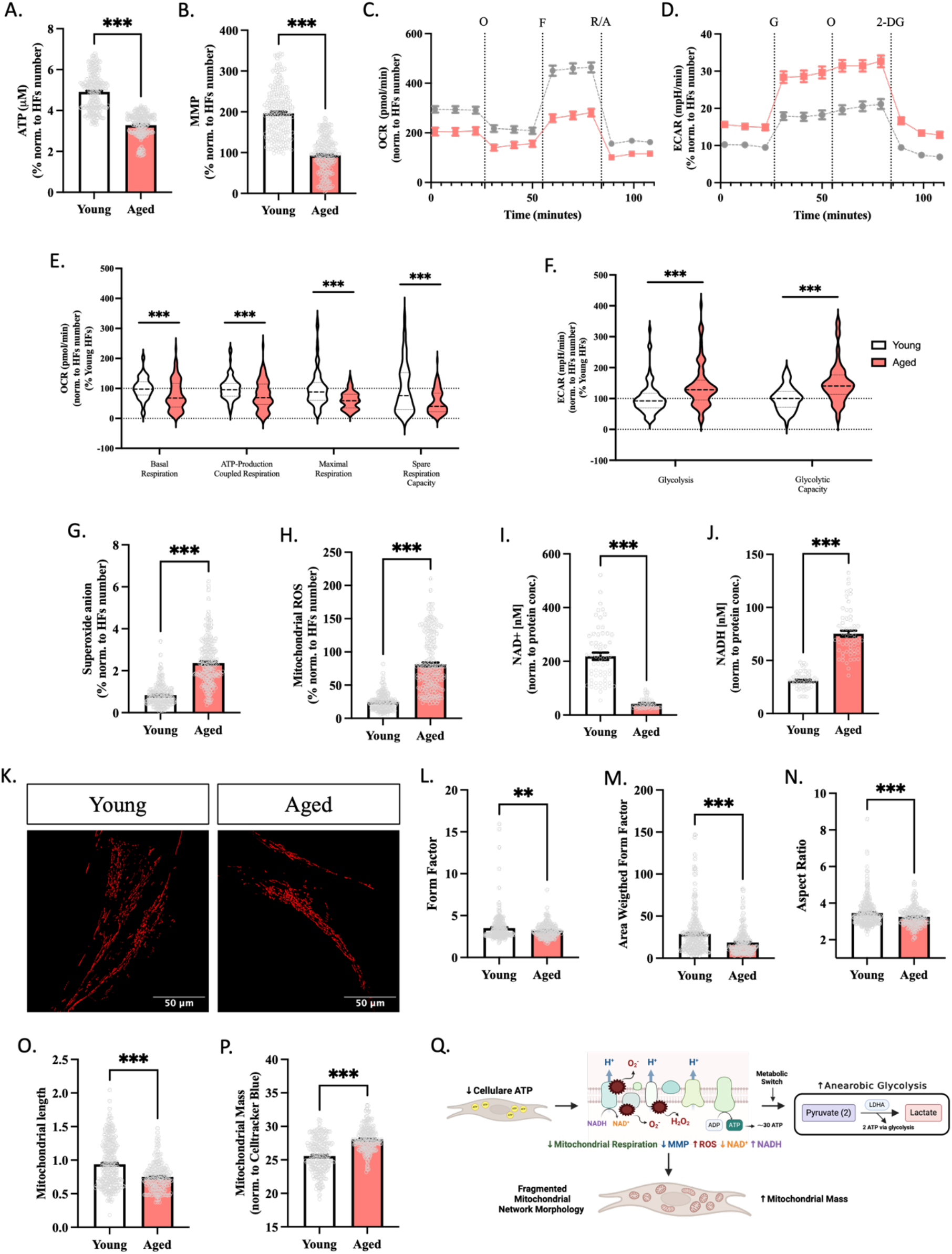
Aging phenotype in aged HFs in comparison of young HFs on mitochondrial properties. A. Cellular ATP level comparing HFs from aged and young donors using a bioluminescence assay. (N= 5 independent experiments, n=13-14 replicates per experiment) B. MMP level comparing aged HFs to young HFs by staining with TMRM. The fluorescence was detected at ex: 548 nm /em: 574 nm. (N= 5 independent experiments, n=10-17 replicates per experiment) C. Mito Stress Test profile representing the OCR of young and aged HFs after sequential injection of oligomycin (O, 1 µM), FCCP (F, 2 µM), and lastly combined rotenone (R, 2 µM) with antimycin A (A, 4 µM). D. Glycolysis Stress Test profile exhibiting the ECAR of aged and young HFs after sequential injection of glucose (G, 10 mM), Oligomycin (O, 1µM), and lastly 2-deoxy-glucose (2-DG, 25 mM). E. Bioenergetic parameters calculated from the Agilent Seahorse XF Cell Mito Stress Test of young and aged HFs. Basal respiration, ATP-production coupled respiration, maximal respiration, and spare respiration capacity. (N= 5-6 independent experiments, n=3-5 replicates per experiment) F. Bioenergetic parameters were determined from the Agilent Seahorse XF Glycolysis Stress Test by comparing young and aged HFs. Glycolysis and glycolytic capacity. (N= 4-6 independent experiments, n=2-4 replicates per experiment) G. Mitochondrial superoxide anione using the MitoSOX dye to compare young and aged HFs. The fluorescence was detected at ex: 485 nm /em: 535 nm. (N= 5 independent experiments, n=11-12 replicates per experiment) H. Mitochondrial ROS detection in aged and young HFs. (ex: 485 nm /em: 535 nm). The fluorescence was detected at ex: 531 nm /em: 595 nm. (N= 5 independent experiments, n=11-12 replicates per experiment) I,J. Cellular NAD+ content (I) and NADH content (J) from young and aged HFs represented as normalised values to the protein concentration. (N= 5 independent experiments, n=3 replicates per experiment) K-O. Mitochondrial network morphology was assessed in HFs from young and aged human donors by staining the mitochondria with TOMM20. Calculated mitochondrial parameters Form Factor (L), Area Weighted Form Factor (M), Aspect Ratio (N), and Length (O). (N= 4-5 independent experiments, n=12-33 replicates per experiment) P. Mitochondrial Mass comparing young and aged HFs assessed by using the MitoTracker™ Green FM (ex: 490 nm /em: 516 nm) to stain mitochondria and were normalized to the cell area using CellTracker™ Blue CMAC Dye (ex: 353 nm /em: 466 nm). (N= 4 independent experiments, n=15-16 replicates per experiment) Q. Schematic representation of the findings regarding the aged HFs phenotypes in comparison to HFs from young donors. The up arrow ↑ represents an increase in the representative parameter and the down arrow ↓ indicates a decrease in the representative parameters. Created with BioRender.com **Data information:** All data are represented as the mean ± SEM of each 4 different young and aged HFs. Values were normalized on the cell count by visualizing the nucleus with DAPI staining, except for the cellular ATP, NAD^+^/ NADH ratio, mitochondrial network morphology, and mitochondrial mass after the experiments. Student’s unpaired t-test was performed for young HFs versus aged HFs (* p < 0.05, ** p < 0.01, *** p < 0.001). Abbreviation: 2-DG: 2 deoxy-glucose; ATP: Adenosine triphosphate; ECAR: extracellular acidification rate; F: FCCP; G: glucose; HFs: Human fibroblasts; Max: maximal; Mito: Mitochondria, MMP: mitochondrial membrane potential; O: oligomycin; OCR: oxygen consumption rate; R: rotenone. Please refer to the data availability section.

Subsequently, the reduction/oxidation (redox) state was quantified by measuring mitochondrial-derived free radicals and reactive oxygen species (ROS) (Fig 1G & H) levels and nicotinamide adenine dinucleotide (NAD) by quantifying NAD^+^ to NADH ratio (Fig 1I & J). While mitochondria are the leading players in cellular energy production, they also serve as the primary source of ROS, mainly generated as unavoidable by-products of ETC [45]. We quantified the levels of mitochondrial superoxide anion radicals (Fig 1G) and mitochondrial ROS (Fig 1H), where we observed a substantial increase in both mitochondrial free radical levels in aged donors. Our investigation on NAD revealed that the concentration of NAD^+^ decreased (Fig 1I) while the concentration of NADH increased (Fig 1J) in aged HFs compared to young HFs. The NAD^+^/NADH ratio decreased from 7.1 in young HFs to 0.56 in aged HFs (EV Table 1). The NAD is a crucial redox molecule involved in cell metabolism and signaling [46, 47]. The intracellular NAD^+^ to NADH redox state can be seen as an indicator of the metabolic balance between the cellular glucose and oxygen metabolisms. Thus, the ratio could function as the oxygen-to-glucose index, a crucial biological indicator of cellular bioenergetics [46].

**Table 1:**
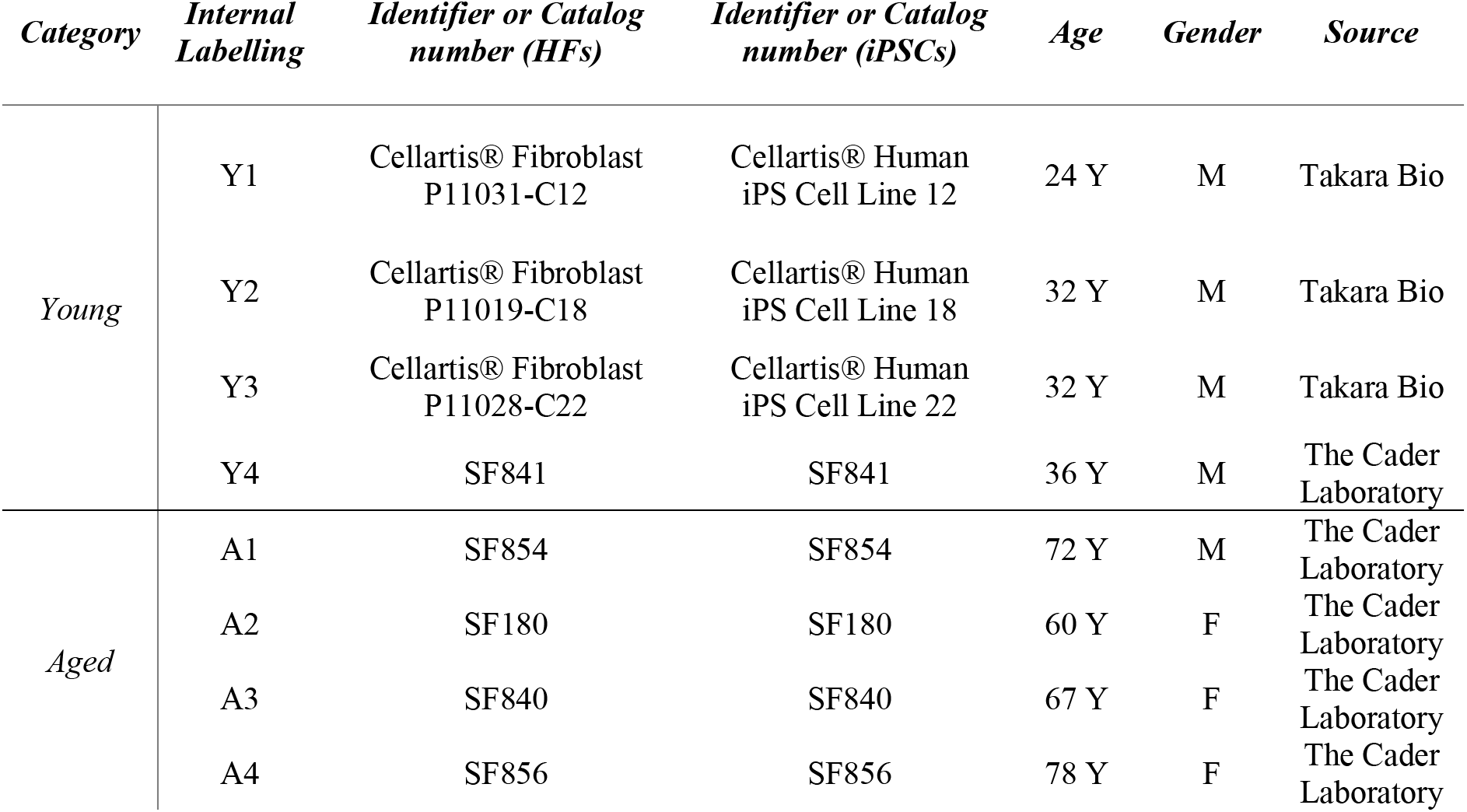
Donor information of the HFs and iPSCs. F: Female; M: Male; Y: Year

| *Category* | *Internal Labelling* | *Identifier or Catalog number (HFs)* | *Identifier or Catalog number (iPSCs)* | *Age* | *Gender* | *Source* |
| --- | --- | --- | --- | --- | --- | --- |
| *Young* | Y1 | Cellartis® Fibroblast P11031-C12 | Cellartis® Human iPS Cell Line 12 | 24 Y | M | Takara Bio |
|  | Y2 | Cellartis® Fibroblast P11019-C18 | Cellartis® Human iPS Cell Line 18 | 32 Y | M | Takara Bio |
|  | Y3 | Cellartis® Fibroblast P11028-C22 | Cellartis® Human iPS Cell Line 22 | 32 Y | M | Takara Bio |
|  | Y4 | SF841 | SF841 | 36 Y | M | The Cader Laboratory |
| *Aged* | A1 | SF854 | SF854 | 72 Y | M | The Cader Laboratory |
|  | A2 | SF180 | SF180 | 60 Y | F | The Cader Laboratory |
|  | A3 | SF840 | SF840 | 67 Y | F | The Cader Laboratory |
|  | A4 | SF856 | SF856 | 78 Y | F | The Cader Laboratory |

Crucial to their bioenergetic status, mitochondria are morphologically dynamic organelles that form an interconnected or fragmented network through fission and fusion events [10, 48–51]. The mitochondrial network morphology was quantified further by visualizing the mitochondria in young and aged HFs using a TOMM20 staining. The mitochondrial network morphology analysis showed significant differences in all calculated parameters between aged and young HFs (Fig 1L-O). We found that aged HFs exhibited a more fragmented mitochondrial state. This was indicated by a significantly lower Form Factor (FF) (Fig 1L), Area Weighted Form Factor (AW) (Fig 1M), and Aspect Ratio (AR) (Fig 1N), which all describe a more circular mitochondrial morphology in aging. Additionally, we measured that aged HFs represented shorter mitochondrial lengths (Fig 1O), further supporting a fragmented mitochondrial network in aged HFs. Upon visual comparison of the mitochondrial network (Fig 1K), young HFs appeared to have a more tubular or elongated morphology. In contrast, aged HFs exhibited a more fragmented mitochondrial network. Our findings on the mitochondrial network morphology hinted toward increased fission-related dynamics to remove dysfunctional mitochondria in age. Next, the mitochondrial mass was evaluated to observe the buildup of malfunctioning mitochondria. In investigating the mitochondrial mass (Fig 1O), we observed an increase in mitochondrial mass in aged HFs compared to younger HFs.

In Fig 1Q, we have summarized the aging characteristics of mitochondrial properties observed in our aged HFs compared to HFs from young donors. With this, we demonstrated a decline in cellular ATP, MMP, mitochondrial respiration, and NAD^+^ to NADH ratio and a rise in anaerobic glycolysis and free radicals. Moreover, we detected a fragmented mitochondrial network morphology and elevated mitochondrial mass in aged HFs, strongly suggesting an accumulation of dysfunctional mitochondria. The initial part of this study aimed to determine the mitochondrial impairment in the donor cells, HFs. Subsequently, we examined the iNs and iPSCsNs derived from the same donor cells. Our goal was to assess to which degree the aging donor characteristics are maintained in these cellular models.

### 4.2. Aged iNs displayed donor-associated deficits in comparison to young iNs

In this comparative analysis between iNs and iPSCsNs, we first assessed whether aged iNs served as a reliable *in vitro* model system for neuronal aging. Foremost, an investigation was conducted on the bioenergetic parameters, revealing a decrease in cellular ATP levels (Fig 2A) in aged iNs when compared to young iNs. Upon examination of the MMP (Fig 2B), a marker indicating the health of mitochondria, a significant decline was observed in aged iNs. Furthermore, the levels of mitochondrial superoxide anion radicals (Fig 2C) and mitochondrial ROS (Fig 2D) were measured to detect the emission of mitochondrial radicals. The results indicate that both parameters were significantly higher in aged iNs than young iNs. In the next step, the mRNA expression of antioxidant defense mechanism-related proteins, namely superoxide dismutase 1 (SOD1), catalase (CAT), and glutathione peroxidase (GPX1), were examined to determine their levels in young and aged iNs (Fig 2E). The findings revealed no statistically significant disparities in the mRNA expression of *SOD1* and *GPX1* between aged and young iNs. Nevertheless, the expression of *CAT* was markedly increased in aged iNs.

**Figure 2:**
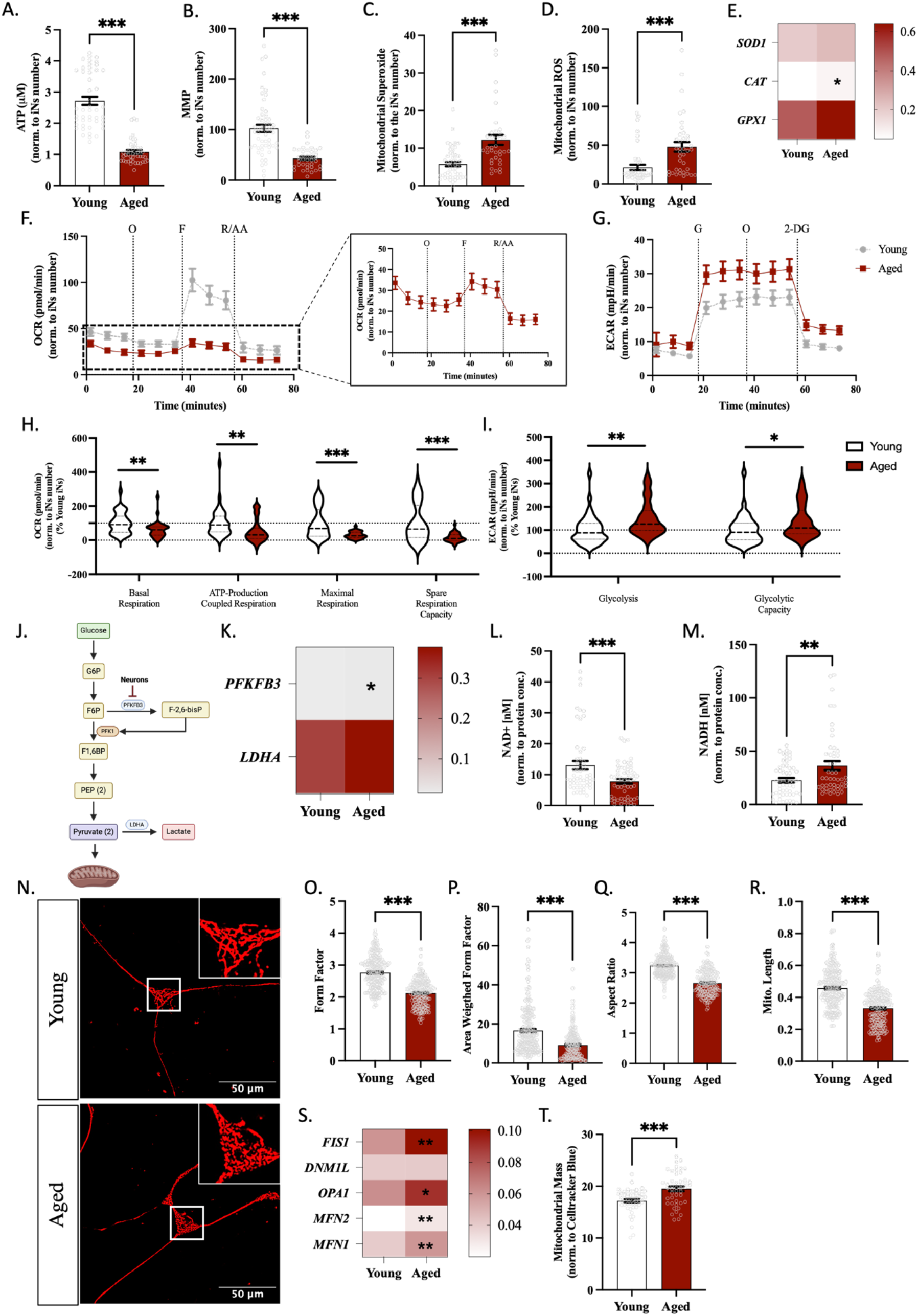
Aged iNs represented aging-associated phenotype from the donor on the mitochondrial level and the correlated metabolic shift toward anaerobic glycolysis. A. Cellular ATP level comparing iNs from aged and young human donors. (N= 3-4 independent experiments, n=2-3 replicates per experiment) B. MMP level measured in aged iNs to young iNs by staining them with TMRM. The fluorescence was detected at ex: 548 nm /em: 574 nm. (N= 4-5 independent experiments, n=2-3 replicates per experiment) C. Mitochondrial superoxide anion detection using the MitoSOX dye to compare young and aged iNs. The fluorescence was detected at ex: 485 nm /em: 535 nm. (N= 4-5 independent experiments, n=2-3 replicates per experiment) D. Mitochondrial ROS detection in aged and young HFs. (ex: 485 nm /em: 535 nm). The fluorescence was detected at ex: 531 nm /em: 595 nm. (N= 4-5 independent experiments, n=2-3 replicates per experiment) E. Relative gene expression of relevant anti-oxidative stress enzymes, *SOD1, CAT*, and *GPX1*. The data are represented as gene expression (2 ^(-Avg.(Delta(Ct))^) as Housekeeping gene *GAPDH* was utilized. F. Mito Stress Test profile representing the OCR of young and aged iNs after sequential injection of oligomycin (O, 2,5 µM), FCCP (F, 2 µM), and lastly combined rotenone (R, 2 µM) with antimycin A (A, 2 µM) G. Glycolysis Stress Test profile representing the ECAR of aged and young iNs after sequential injection of glucose (G, 10 mM), Oligomycin (O, 1µM), and lastly, 2-deoxy-glucose (2-DG, 25 mM) H. Bioenergetic parameters of the mitochondria of young and aged iNs. Basal respiration, ATP-production coupled respiration, proton leak, maximal respiration, and spare respiration capacity. (N= 4-5 independent experiments, n=2-3 replicates per experiment) I. Bioenergetic parameters of glycolysis comparing young and aged iNs. Glycolysis, glycolytic capacity, and glycolytic reverse. (N= 3-5 independent experiments, n=2-3 replicates per experiment) J. Simplified representation of glycolysis in the neurons for illustrative purposes. Created with BioRender.com K. Relative gene expression of relevant glycolysis regulating genes, *PFKFB3* and *LDHA*. The data are represented as gene expression (2 ^(-Avg.(Delta(Ct))^) as Housekeeping gene *GAPDH* was utilized. L,M. Cellular NAD^+^ content (L) and NADH (M) content from young and aged iNs represented as normalized values to the protein concentration. (N= 4-5 independent experiments, n=2-3 replicates per experiment) N-R. The mitochondrial network morphology (N) visualization was assessed in iNs from young and aged human donors by staining the mitochondria with TOMM20. Calculated mitochondrial parameters Form Factor (O), Area Weighted Form Factor (P), Aspect Ratio (Q), and Length (R). (N= 4-5 independent experiments, n=12-33 replicates per experiment) S. Relative gene expression of relevant genes involved in mitochondrial dynamics: *FIS1, DNM1L, OPA1*, *MFN2*, and *MFN1*. The data are represented as gene expression (2 ^(-Avg.(Delta(Ct))^) as Housekeeping gene *GAPDH* was utilized. T. Mitochondrial Mass comparing young and aged iNs assessed by using the MitoTracker™ Green FM (ex: 490 nm /em: 516 nm) to stain mitochondria and were normalized to the cell area using Celltracker blue (ex: 353 nm /em: 466 nm). (N= 4-5 independent experiments, n=2-3 replicates per experiment) **Data information:** All data are represented as the mean ± SEM of each 4 different young and aged iNs. Only three donors were assessed for the gene expression using three technical replicates. Values were normalized on the cell count by visualizing the nucleus with DAPI staining, except for the ATP, NAD^+^/ NADH ratio, mitochondrial network morphology, and mitochondrial mass after the experiments. The representative images were choosen for visualization purposes. Student’s unpaired t-test was performed for young iNs versus aged iNs (* p < 0.05, ** p < 0.01, *** p < 0.001). Abbreviation: 2-DG: 2 deoxy-glucose; ATP: Adenosine triphosphate; CAT: Catalase; DNM1L: Dynamin-related protein 1 (Drp1); ECAR: extracellular acidification rate; F: FCCP; FIS1: Mitochondrial Fission Protein 1; F1,6BP: Fructose 1,6-bisphosphate; F2,6BP: fructose-2,6-bisphosphate; F6P: Fructose-6-phopshate; G: glucose; GPX1: Glutathione peroxidase 1; G6P: Glucose-6-phophaste; iNs: directly converted neurons; LDHA: Lactate dehydrogenase A (LDHA); Max: maximal; MFN1: Mitofusin-1; MFN2: Mitofusin-2; Mito: Mitochondria, MMP: mitochondrial membrane potential; PFKFB3: 6-phosphofructo-2-kinase/fructose-2,6-biphosphatase 3; PEP: phophoenolpyruvic acid: PFK-1:Phosphofructokinase-1; O: oligomycin; OCR: oxygen consumption rate; R: rotenone; Superoxide dismutase 1;. Please refer to the data availability section.

Afterward, the Seahorse XF Cell Mito Stress Test Kit (Fig 2F) was conducted to obtain a thorough evaluation of mitochondrial respiration. A significant decrease in the calculated mitochondrial bioenergetic parameters was observed when comparing young and aged iNs, as shown in Fig 2H. Furthermore, the Seahorse Glycolysis Stress Test (Fig 2G) was conducted to evaluate the likelihood of a metabolic shift towards anaerobic glycolysis, a phenomenon linked to aging. Upon examining the parameters of the Glycolysis Stress Test (Fig 2D), we observed that aged iNs demonstrated a notable elevation in both glycolysis and glycolytic capacity compared to young iNs. This suggests that the shift towards a more anaerobic glycolytic metabolism with aging persisted in the aged iNs after direct conversion. Our study aimed to delve deeper into the mechanism of glycolysis [52]. To gain valuable insights into this intricate system, we analyzed the gene expression of two critical enzymes that play a crucial role in glycolysis, 6-phosphofructo-2-kinase/fructose-2, 6-bisphosphatase-3 (PFKFB3) and lactate dehydrogenase A (*LDHA*). Fig 2J illustrates a simplified diagram of the glycolytic pathway. While studying the *PFKF3B* and *LDHA* gene expression (Fig 2K), we observed that the expression of *PFKF3B* was significantly upregulated in aged iNs. In contrast, the expression of *LDHA* showed no significant differences in the relative mRNA expression.

Subsequently, the redox state of NAD^+^ to NADH was determined. Upon conducting the analysis of the NAD^+^/NADH redox state, we observed that aged iNs exhibited a significant decline in NAD^+^ levels (Fig 2L) and a simultaneous increase in NADH levels (Fig 2M). This resulted in a significant drop in the ratio of NAD^+^/NADH, decreasing from 0.53 in young iNs to 0.22 in the aged iNs (EV Table 1). As previously described, it was observed that aged HFs had a more fragmented mitochondrial network structure, which was associated with a decline in mitochondrial activity compared to younger HFs.

While examining the organization of the mitochondrial network, we observed that aged iNs exhibited a more fragmented mitochondrial morphology compared to iNs from young donors. A decrease in FF (Fig 2O) was observed alongside a reduction in AW (Fig 2P) and AR (Fig 2Q), suggesting a more round and circular mitochondrial morphology in aged iNs compared to young iNs. In addition, the mitochondria (Fig 2R) in aged iNs exhibit reduced length compared to those in younger iNs, indicating that younger iNs possess a more elongated morphology. After conducting a visual analysis (Fig 2N) of young and aged iNs, a noticeable fragmentation was observed with age. By analyzing the relative gene expression of proteins involved in mitochondrial dynamics, we detected an increase in the expression of genes responsible for both mitochondrial fusion and fission (Fig 2S). More precisely, we observed a significant upregulation in the expression of the gene *FIS1*, which is responsible for fission, and *MFN1*, *MFN2*, and *OPA1*, which are crucial for fusion, in aged iNs compared to young ones. Only the gene *DNML1* encoding for DRP1 did not indicate differences in the gene expression. Additionally, an elevated mitochondrial mass (Fig 2T) was observed in aged iNs compared to young iNs. The phenotype of the aged iNs closely resembled that of the aged HFs, suggesting that the iNs displayed the characteristic traits of aging as observed in the donor cells.

### 4.3. Aged iPSCsNs preserved donor aging signature on the mitochondrial state except not on the glycolytic level

We conducted the same measurements in the iPSCsNs derived from the same donors as in iNs. Initially, we measured the cellular ATP level (Fig 3A) in iPSCsNs. Our observations indicate a significant decline in the aged iPSCsNs compared to the young ones in the ATP level. Furthermore, we also found that the MMP (Fig 3B) was decreased. The superoxide anion radical produced by mitochondria (Fig 3C) and the mitochondrial ROS (shown in Fig 3D) demonstrated aging traits by displaying a notable rise in aged iPSCsNs compared to young iPSCsNs. After investigating the gene expression that plays a vital role in the antioxidant defense mechanism against ROS (Fig 3E), we only found a significant increase in *GPX1* mRNA expression in aged iPSCsNs.

**Figure 3:**
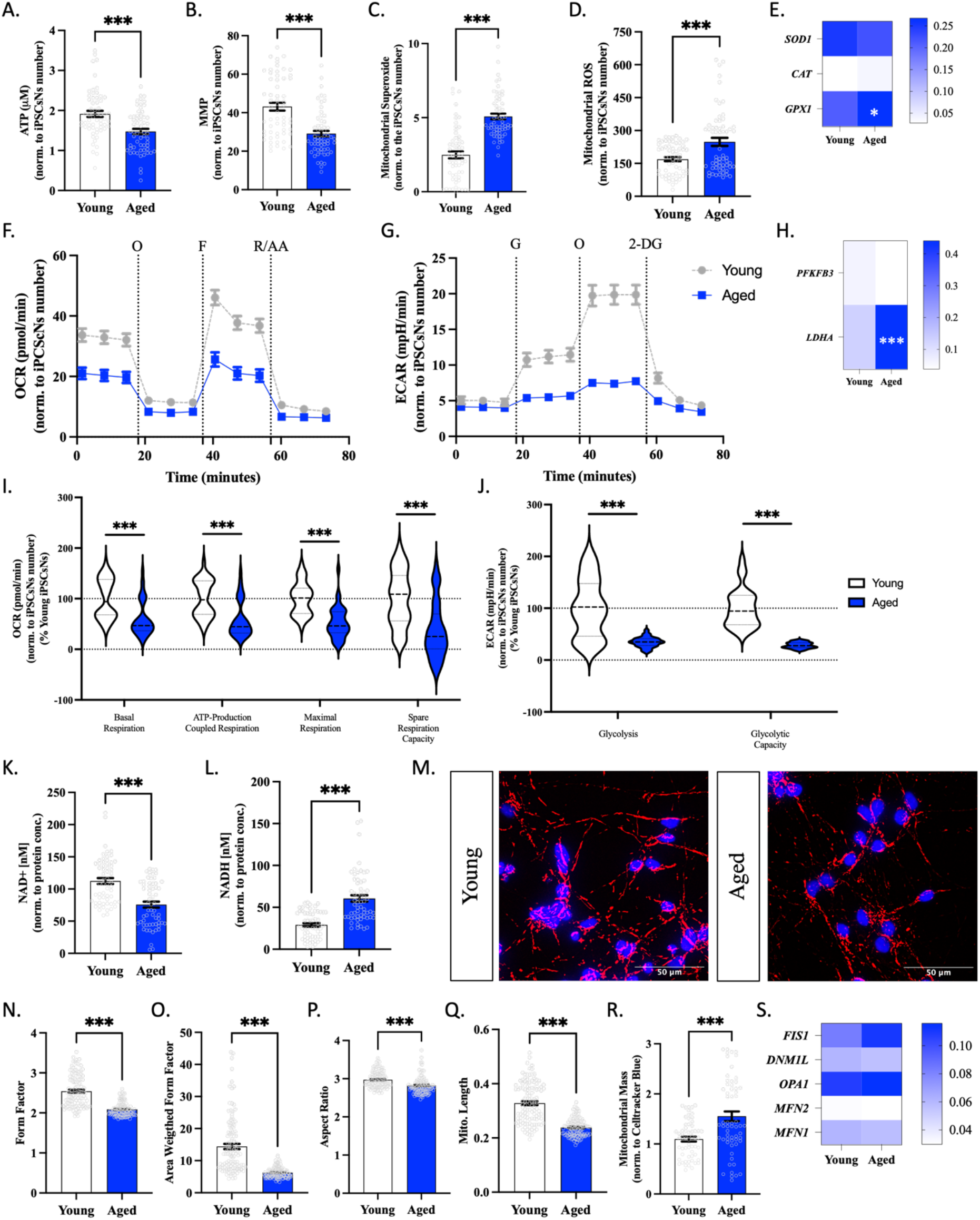
A decrease in mitochondrial function was seen in aged iPSCsNs compared to young iPSCsNs; however, there was no evidence of a metabolic shift towards anaerobic glycolysis. A. Cellular ATP level comparing iPSCsNs from aged and young human donors. (N= 5 independent experiments, n=2-3 replicates per experiment) B. MMP level measured in aged iPSCsNs to young iPSCsNs by staining with TMRM. The fluorescence was detected at ex: 548 nm /em: 574 nm. (N= 5 independent experiments, n=2-3 replicates per experiment) C. Mitochondrial superoxide anion detection using the MitoSOX dye to compare young and aged iPSCsNs. The fluorescence was detected at ex: 485 nm /em: 535 nm. (N= 5 independent experiments, n=3 replicates per experiment) D. Mitochondrial ROS detection in aged and young iPSCsNs. (ex: 485 nm /em: 535 nm). The fluorescence was detected at ex: 531 nm /em: 595 nm. (N= 5 independent experiments, n=3 replicates per experiment) E. Relative gene expression of relevant anti-oxidative stress enzymes, *SOD1, CAT*, and *GPX1*. The data are represented as gene expression (2 ^(-Avg.(Delta(Ct))^) as Housekeeping gene GAPDH was utilized. F. Mito Stress Test profile representing the OCR of young and aged iPSCsNs after sequential injection of oligomycin (O, 2,5 µM), FCCP (F, 2 µM), and lastly combined rotenone (R, 2 µM) with antimycin A (A, 2 µM). G. Glycolysis Stress Test profile representing the ECAR of aged and young iPSCsNs after sequential injection of glucose (G, 10 mM), Oligomycin (O, 1µM), and lastly 2-deoxy-glucose (2-DG, 25 mM). H. Bioenergetic parameters of the mitochondria of young and aged iPSCsNs. Basal respiration, ATP-production coupled respiration, maximal respiration, and spare respiration capacity. (N= 4 independent experiments, n=2-3 replicates per experiment) I. Bioenergetic parameters of glycolysis comparing young and aged iPSCsNs. Glycolysis and glycolytic capacity. (N= 3 independent experiments, n=2-3 replicates per experiment) J. Relative gene expression of relevant glycolysis-regulating genes, *PFKFB3, PKM, LDHA*. The data are represented as gene expression (2 ^(-Avg.(Delta(Ct))^) as the Housekeeping gene GAPDH was utilized. K,L. Cellular NAD^+^ content (K) and NADH content (L) from young and aged iPSCsNs represented as normalized values to the protein concentration. (N= 5 independent experiments, n=3 replicates per experiment) M-Q. Mitochondrial network morphology (M) was assessed in iPSCsNs from young and aged human donors by visualizing the mitochondria with TOMM20 and nucleus staining with DAPI. Calculated mitochondrial parameters Form Factor (N), Area Weighted Form Factor (O), Aspect Ratio (P), and Length (Q). (N= 5 independent experiments, n=3 replicates per experiment). R. Mitochondrial Mass comparing young and aged iPSCsNs assessed by using the MitoTracker™ Green FM (ex: 490 nm /em: 516 nm) to stain mitochondria and were normalized to the cell area using Celltracker blue (ex: 353 nm /em: 466 nm). (N= 5 independent experiments, n=3 replicates per experiment) S. Relative gene expression of relevant genes involved in mitochondrial dynamics: *FIS1, DNM1L, OPA1, MFN2*, and *MFN1*. The data are represented as gene expression (2 ^(-Avg.(Delta(Ct))^) as the Housekeeping gene GAPDH was utilized. **Data information:** All data are represented as the mean ± SEM of each 4 different young and aged iNs. Only three donors were assessed for the gene expression with three technical replicates. Values were normalized on the cell count by visualizing the nucleus with DAPI staining, except for the ATP, NAD^+^/ NADH ratio, mitochondrial network morphology, and mitochondrial mass after the experiments. The representative images were produced for visualization purposes. Student’s unpaired t-test was performed for young iNs versus aged iNs (* p < 0.05, ** p < 0.01, *** p < 0.001). Abbreviation: 2-DG: 2 deoxy-glucose; ATP: Adenosine triphosphate; CAT: Catalase; DNM1L: Dynamin-related protein 1 (Drp1); ECAR: extracellular acidification rate; F: FCCP; FIS1: Mitochondrial Fission Protein 1; G: glucose; GPX1: Glutathione peroxidase 1; iPSCsNs: induced pluripotent stem cells (iPSCs) derived neurons; LDHA: Lactate dehydrogenase A (LDHA); Max: maximal; MFN1: Mitofusin-1; MFN2: Mitofusin-2; Mito: Mitochondria, MMP: mitochondrial membrane potential; PFKFB3: 6-phosphofructo-2-kinase/fructose-2,6-biphosphatase 3; O: oligomycin; OCR: oxygen consumption rate; R: rotenone; Superoxide dismutase 1. Please refer to the data availability section.

During our investigation, we monitored the mitochondrial respiration in real-time using the Seahorse Mito Stress Test (Fig 3F). By analyzing the calculated parameters (Fig 3I) from the OCR graph, we discovered an overall significant decrease in the mitochondrial respiration capacity. After conducting the Seahorse Glycolysis Stress Test (Fig 3G), we identified a significant decline in the glycolysis parameter (Fig 3J) in the aged iPSCsNs. This stands in contrast to the aging phenotype observed in the aged HFs and aged iNs. On analyzing the gene expression level of aged iPSCsNs, it was found that the critical gene on glycolysis, *PFKF3B*, remained unchanged. Nevertheless, it was observed that the levels of *LDHA* mRNA expression in aged iPSCsNs were more significant when compared to young iPSCsNs, indicating a notable difference in LDHA activity between the two groups.

Upon evaluating the NAD^+^ to NADH ratio, we observed a substantial decrease in the NAD^+^ level (Fig 3K). In contrast, a significant increase in the NADH level (Fig 3L) was noted. The NAD^+^/ NADH redox ratio (EV Table 1) was calculated to be 3.87 in young iPSCsNs. In contrast, it was found that the ratio was 1.25 in aged iPSCsNs, indicating a substantial drop in the NAD^+^/NADH redox ratio within the aged iPSCsNs to young iPSCsNs.

While examining the structures of the mitochondrial network morphology in iPSCsNs, we observed that the aged iPSCsNs exhibited a more fragmented mitochondrial network morphology than the younger iPSCsNs. An evident disparity in aged iPSCsNs was noted through a visual comparison (Fig 2M) of young and aged iPSCsNs. The visual assessment of aged iPSCsNs revealed a more fragmented mitochondrial network morphology, as evidenced by a drop in FF (Fig 3N), AW (Fig 3O), and AR (Fig 3P). In addition, the mitochondria length (Fig 3Q) in aged iPSCsNs exhibited a reduced length compared to those observed in younger iPSCsNs, indicating that younger iPSCsNs possessed a more extended morphology. Our study observed that the mitochondrial mass (Fig 3Q) was higher in aged iPSCsNs than in young iPSCsNs. However, no significant changes were found in the expression of genes associated with mitochondrial dynamics between young and aged iPSCsNs.

### 4.4. Transcriptomic alteration in the neuronal models of aging, iNs and iPSCsNs, of genes related to mitochondrial properties

Previous research has shown that reprogramming to the pluripotent state resulted in resetting of aging-associated gene expression [23]. Only directly converted neurons could preserve the aging-associated donor signature on the transcriptomic level. With that objective in mind, we investigated the mRNA expression of further crucial genes related to mitochondrial characteristics (Fig 4A). Our investigation revealed notable alterations in the mRNA expression of poly (ADP-ribose)-polymerase-1 (PARP1), Serine/threonine-protein kinases 1 (AKT1), AMP-activated protein kinase (AMPK), Nuclear factor erythroid-derived 2-like 2 (NRF2), Forkhead box protein O1 (FOXO1), and uncoupling protein 2 (UCP2) in aged iNs when compared to young iNs, in addition to the previously identified genes. The mRNA expression of NRF2, FOXO1, and UCP2 in aged iPSCsNs differed significantly from that in young iPSCsNs. Nevertheless, the remaining genes exhibited no alteration. The following section provides a summary of the main results of our transcriptomic analysis. Along with NRF2 and mitochondrial transcription factor A (TFAM), peroxisome proliferator-activated receptor gamma coactivator 1 alpha (PGC-1α) triggers mitochondrial biogenesis (mitogenesis) [51, 53]. Regarding the regulation of these genes for mitogenesis, we found that only the *NRF2* gene showed a notable increase in expression in both aged iNs and aged iPSCsNs. Upon examining the upstream regulator of PGC-1α, we discovered that *FOXO1*, a forkhead transcription factor, was significantly downregulated in aged iNs and upregulated in aged iPSCs in direct comparison to the corresponding young. Another upstream regulator, AMPK, showed to be elevated in aged iNs, but no significant disparities in AMPK gene expression were observed between aged and young iPSCsNs [51]. Further investigation was conducted into the inhibitory pathways of PGC-1α, including AKT1, a mTOR activator [51]. In turn, mTOR is an inhibitor of the autophagy/mitophagy. For the *AKT1* mRNA expression, only aged iNs showed a significant upregulation than young iNs, while there was no discernible difference between the aged and young iPSCsNs. Other genes that showed a significant upregulation in the aged iNs, as reflected in aging [54], but not in the aged iPSCsNs to the corresponding young, were *PARP1* and *UCP2*. The aged iNs, in direct comparison to young iNs, showed a significant rise in expression, whereas the aged iPSCsNs showed a significant downregulation than young iPSCsNs. Given the more pronounced changes observed in the aged iNs compared to aged iPSCs, we investigated whether there would be differences in the mRNA expression levels between these two neuronal models. Therefore, we performed a principal component analysis (PCA) comparing the gene expression profile of all detected genes of aged iNs versus aged iPSCsNs against each other. Using the first two principal components to plot our data set, we visually identified that the data set of aged iNs and aged iPSCsNs did not cluster (Fig 4B).

**Figure 4:**
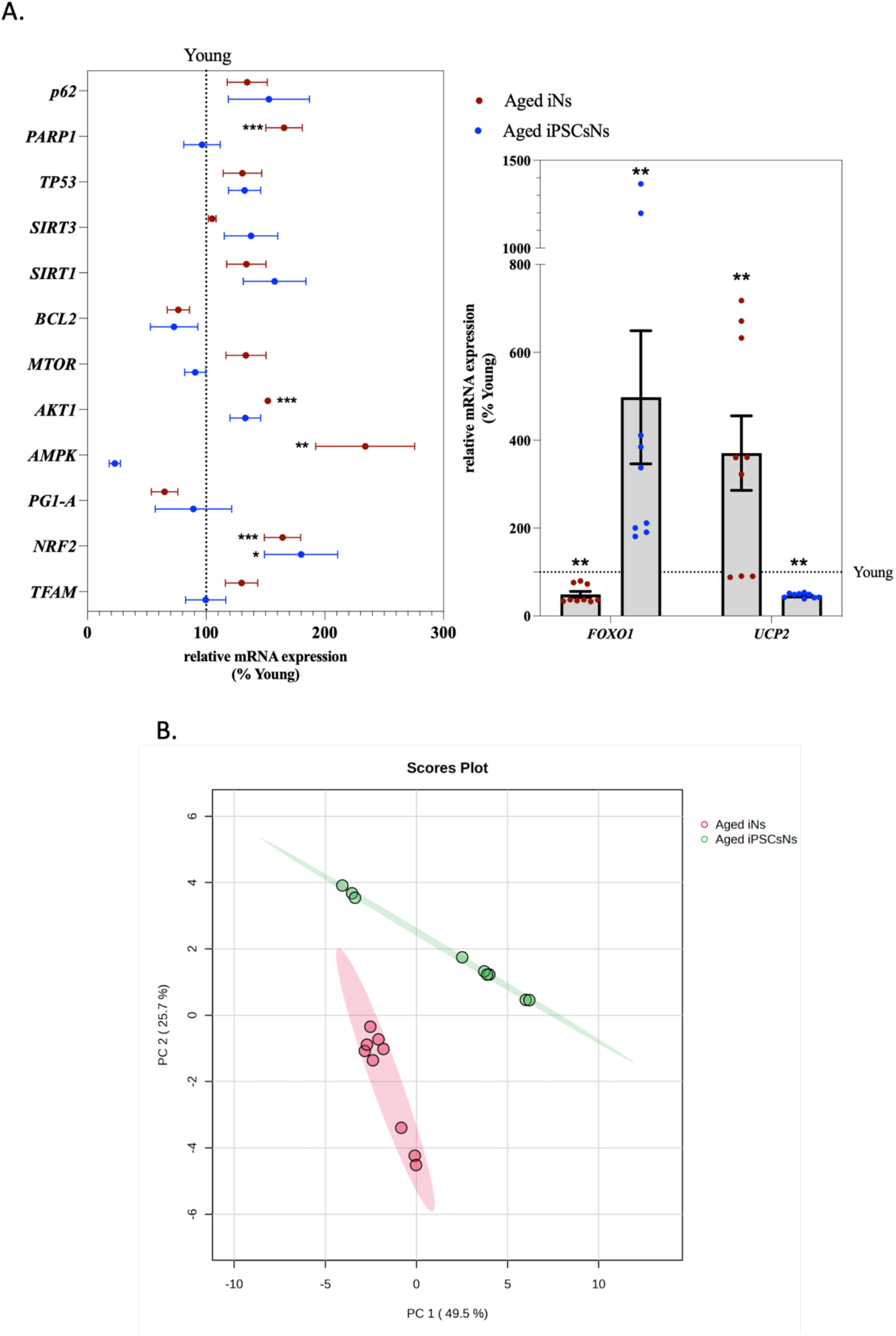
Genes related to mitochondrial properties show transcriptomic alterations in neuronal models of aging. A. The mRNA expression profile of aged iNs and aged iPSCsNs % normalized to the corresponding young. The smaller graph represented the mRNA expression of the genes *FOXO1* and *UCP2*, which had the highest percentage difference compared to the corresponding young. (N= 3 independent experiments, n=3 replicates per experiment) B. Aged iNs *versus* aged iPSCsNs of detected genes in our investigation. Principal component analysis of aged iNs (red) and aged iPSCsNs (green). Scores plot between the selected PC1 and PC2. Each collared dot represents a replicate. (N= 3 independent experiments, n=3 replicates per experiment) **Data information:** The represented values for the gene expression were assessed from 3 donors with 3 technical replicates for each condition. Student’s unpaired t-test was performed for young versus aged (* p < 0.05, ** p < 0.01, *** p < 0.001). Figure B was generated on metaboanalyst.ca. The represented values were normalized by autoscaling (mean-centered and divided by the standard deviation of each variable). The data were represented as Gene expression (2^(-Avg.(Delta(Ct))^) by using the Housekeeping gene *GAPDH* calculated by the website https://geneglobe.qiagen.com/us/analyze. Figure B was generated using <u>metaboanalyst.ca</u>. Please refer to the data availability section. Abbreviation: AKT1: Serine/threonine-protein kinases 1; BCL2: B-cell lymphoma 2; FOXO1: Forkhead box protein O1; HEK293: Human Embryonic Kidney 293; HFs: Human fibroblasts; iNs: Induced neurons; iPSCs: induced pluripotent stem cells; iPSCsNs: iPSCs MFN1: Mitofusin-1; MFN2: Mitofusin-2; Mito: Mitochondria; MMP: mitochondrial membrane potential; MTOR: Mammalian target of rapamycin; NFE2L2: Nuclear factor erythroid-derived 2-like 2 (NRF2); OPA1: Optic atrophy 1; PARP1: Poly (ADP-ribose) polymerase 1; PFKFB3: 6-phosphofructo-2-kinase/fructose-2,6-biphosphatase 3; PPARGC1A: Peroxisome proliferator-activated receptor gamma coactivator 1-alpha (PGC-1α); PRKAA2: AMP-activated protein kinase (AMPK); SIRT1: Silent mating type information regulation 2 homolog 1; SIRT3: Silent mating type information regulation 2 homolog 3; SQSTM1:Sequestosome 1 (p62); TFAM: Mitochondrial transcription factor; TP53: Transformation-related protein 53 (p53); UCP2: Mitochondrial uncoupling protein 2. Please refer to the data availability section.

**Figure 5:**
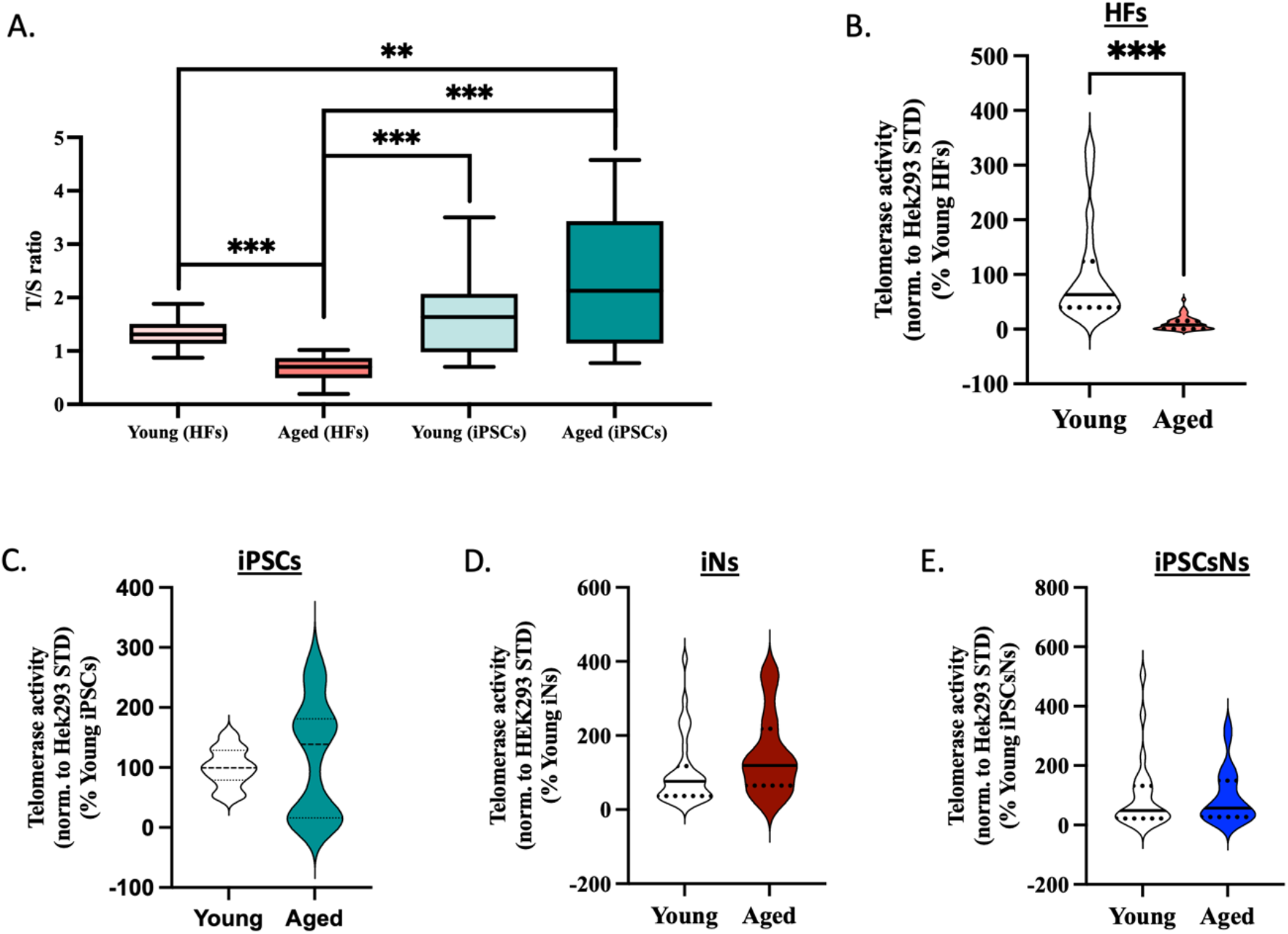
Telomere impairments in aged HFs but not in the derived cell types. A. Telomere length measurement (A) of young and aged HFs and young and aged iPSCs. The data are represented as the TELO/ SGC (T/S). A mixture of all HFs DNA samples was used as an internal control. (N= 4-5 independent experiments, n= 3 replicates per experiment) B-E. The telomerase activity from young and aged donors was measured in HFs (B), iPSCs (C), iNs (D), and iPSCsNs (E). The assessment was done in 10’000 cells for HFs and iPSCs and 100’000 cells for both neuronal models. The data are represented as values normalized to a Hek293 STD and normalized to 100 % of the corresponding young. (N=2-6 independent experiments, n=1-2 replicates per experiment) **Data information:** All data are represented as the mean ± SEM of each 4 different young and aged HFs, iPSCs, iNs, or iPSCsNs. Except for T/S iPSCs, only 3 donors were utilized. The represented values from four young and aged donors derived iNs show N independent experiments with n technical replicates per donor. Student’s unpaired t-test was performed for young versus aged (* p < 0.05, ** p < 0.01, *** p < 0.001). Abbreviation: HFs: Human fibroblasts; iNs: directly induced neurons; iPSCsNs: induced pluripotent stem cells (iPSCs) derived neurons; norm.: normalized. Please refer to the data availability section.

### 4.5. Comparative analysis of mitochondrial parameters on individual donors to understand the plasticity of aging

We conducted a comparative analysis of the critical mitochondrial parameters to understand how the conversions affect the mitochondrial properties at the individual donor level. In a spiderweb comparison (EV Fig 2 and 3) of the various cell models, including HFs, neuronal cell iNs, and iPSCsNs, for each donor, the normalized values were represented in relation to the representative young condition (EV Table 3). Overall, we noticed a significant alignment of the distinct characteristics that were maintained during the transformation process. However, there were noticeable variations in specific characteristics among the individual donors. In our observation of the young donors (EV Fig 2), we noticed notable differences in the maximal respiration capacities among the various cell types. Specifically, when comparing donors Young 2 and Young 3, we found that Young 2 had the highest level of maximal respiratory capacity in the iNs. In contrast, Young 3 experienced a significant decline in the iNs’ state of maximal capacity. Interestingly, the other cell states, HFs and iPSCsNs, did not exhibit such differences. We detected a substantial rise in the mitochondrial ROS parameters and NAD^+^ levels in the iNs state of Young 3 compared to the HFs and iPSCsNs state. We noted that Young 3 showed the most substantial variation between the young donors in the iNs state, while the HFs’ and iPSCsNs’ states appeared more similar to the other donors. In the comparison between the different aged donors (EV Fig3), it was observed that aged donors in different reprogramming states shared some similarities. However, one noticeable difference stood out when comparing the donor phenotype across different age groups. As previously stated, the radical emission in the aged HFs state was more substantial compared to the neuronal state, except for donor Aged 3, which showed for the HFs state low ROS emission compared to the other cell states and donors. All donors showed a glycolysis deficiency in the iPSCsNs state, indicating no metabolic transition from mitochondrial respiration to glycolysis. No additional inconsistencies were observed upon visual examination of the radar plots.

### 4.6. Decrease of telomerase activity was not observed in both aged neuronal models

In our analysis, we further concentrated on telomere length as an additional aging parameter. Shorter telomeres have been linked to accelerated aging and genomic instability [55]. In contrast, longer telomeres can extend cellular lifespan, suggesting that telomere biology is essential to the aging mechanism [56, 57]. Our study showed that TL decreased with age in HFs but remained unchanged between young and aged iPSCs (Fig 6A). Interestingly, we also observed an increase in TL from HFs to iPSCs, with young donors increasing from 1.34 T/S to 1.678 T/S and aged donors increasing from 0.66 T/S to 2.32 T/S after conversion. Unfortunately, we could not extract sufficient DNA from the neuronal cell, which made it challenging to determine the TL. As a result, we shifted our focus to investigate the telomerase activity (TA), which counteracts the telomere attrition in healthy cells [58–61]. With age, it is considered that the telomerase activity declines, leading to the manifestation of the aging phenotype [62]. First, we investigated the TA (Fig 6B) in HFs and found a significant decrease in TA in aged HFs compared to young HFs, indicating an aging-associated phenotype on the telomeric level. However, (Fig 6C) we found no significant differences in TA between young and aged iPSCs. Subsequently, we shifted our focus towards evaluating the TA in aged iNs and iPSCsNs to assess telomere attrition. To detect the activity of neuronal cells, we used a higher number of cells than for HFs and iPSCs, from a cell count of 10’000 to 100’000 cells. Interestingly, our results for iNs (Fig 6D) showed no decline in TA associated with aging, unlike HFs. Furthermore, we also examined the TA in iPSCsNs, as shown in Fig 4H. Our results indicated no significant differences in TA between young and aged iPSCsNs (Fig 6E).

## 5. Discussion

### 5.1. Age-related mitochondrial impairment was retained in the aged iNs and aged iPSCsNs

Investigating the predisposing molecular factors of aging is crucial to promote healthy brain aging and enhance our knowledge of neurodegenerative diseases. In this paper, we compared the two advanced human neuronal models in a dish, iNs and iPSCsNs, derived from the same donor HFs on the aspect of aging. The goal was to determine the most suitable neuronal model organism for studying neuronal aging in vitro. We defined the aging signature of the corresponding donor HFs by comparing young HFs (24 – 36 years) and aged HFs (60 – 78 years) and then the corresponding iNs and iPSCsNs from the same donors. Our research has revealed considerable variations in the mitochondrial bioenergetic status of young and aged HFs, which was observed in the iNs. It is worth noting that the differences were noticed even in the iPSCsNs, which was an unexpected outcome based on previous research [23, 25, 27, 29, 63–65]. The extent to which reprogramming iPSCs and the resulting cells can fully reset the metabolic and cellular alteration associated with aging is currently under debate. While certain studies propose the feasibility of an overall rejuvenation, others indicate that only partial rejuvenation occurs [40]. In our previous study (unpublished data of the Eckert group), we could demonstrate an aging-associated impairment in the aged iPSCs compared to young iPSCs, which we could replicate further in the neuronal state of the same donors. We have shown that aged iPSCsNs exhibit a similar impairment on mitochondrial properties as aged iNs and HFs. This discovery presents new possibilities for creating neuronal *in vitro* models of aging.

The brain is a high-energy consumer, requiring 20 % of the body’s basal oxygen and consuming 25 % of the available glucose to function correctly [66, 67]. Cells require a continuous energy supply to generate and maintain the biological structure essential for their survival [68, 69]. Our first investigation indicated that the overall energy state (ATP level) was lower in the aged HFs, which was also observed in the aged neuronal cells, iNs and iPSCsNs. These results highlight that the aged neurons converted directly or via the iPSCs state exhibited aging-related energy impairments, consistent with previous studies on neurons [70–72]. In healthy neurons, mitochondrial biogenetics provides 90 % of the required energy by oxidizing sugars, lipids, and proteins [48]. As a consequence, mitochondrial dysfunction can severely impact neuronal properties [73]. As mitochondria impairments represent a principal target in neuronal aging [10, 42, 74–76], our investigation focused mainly on mitochondrial properties. Hereby, an apparent decline in mitochondrial bioenergetic activity was indicated by assessing a reduction in mitochondrial respiration parameters and a decline in MMP in our aged HFs. For the iNs and iPSCsNs, we could show that the aged neurons, compared to the young condition, preserved their aging-associated donor signature of HFs after the different reprogramming processes. Our research exhibited that our aged neuronal cells, iNs and iPSCsNs, experience a reduction in energy production due to a decline in the activity of the mitochondrial respiratory chain. This decline in MMP and mitochondrial energy production was consistent with previous research [8, 11, 76, 77].

Additionally, as organisms age, their ability to withstand stress, damage, and disease decreases [78]. The exact causes of this gradual decline still need to be fully understood. However, the oxidative stress theory is widely acknowledged as one of the leading explanations for the aging process [79–81]. According to this theory, free radicals generated during cellular respiration can damage essential cellular components like lipids, proteins, and DNA, leading to aging. The literature suggests that the primary reason for a rise in free radicals is the diminished antioxidant defense mechanism against oxidative stress [10, 74]. Due to its close proximity and lower repair mechanism than nDNA, mtDNA is directly exposed to the toxic environment, leading to the accumulation of mutated mtDNA [10, 49, 74, 82, 83]. These mutations in mtDNA, which mainly encodes genes for mitochondrial respiration, adversely affect the energy production capability of the mitochondria. This impairment results in increased emission of ROS, exacerbating cellular damage and aging [84, 85]. The increase in oxidative stress impairs respiratory activity, induces a vicious cycle of mitochondrial impairments, and eventually leads to cell death [86]. Our findings indicated that the concentration of free radicals, mitochondrial superoxide anion and mitochondrial ROS, were higher in the aged iNs and iPSCsNs than their younger counterparts resembling the aged HFs. ROS is mainly produced as a harmful byproduct of multiple cellular pathways that involve redox reactions, with the most significant contributors being the energy pathways facilitated by the coenzymes NAD [87, 88]. NAD is essential for various metabolic functions, including energy production and DNA integrity maintenance [89, 90]. During glycolysis, beta-oxidation, and the TCA cycle, NADH is produced, which is then utilized by NAD^+^ in the ETC of OXPHOS to generate ATP [91]. When all the energy pathways are functioning correctly, the ratio of NAD^+^ to NADH should be high [92]. A decrease in this ratio strongly indicates cellular and mitochondrial functional impairment [93]. Our findings on the HFs, iNs, and iPSCsNs of aged individuals were consistent with previous studies that suggest a reduction in NAD^+^ levels and a simultaneous increase in NADH levels as individuals age, resulting in a decline in the ratio of NAD^+^ to NADH [70, 84, 94–96].

The evaluation of the morphology of the mitochondrial network revealed further information about the bioenergetic state and overall health of the aged iNs and aged iPSCsNs [49, 83, 97]. The morphological network regulates the localization, morphology, and dimensions of mitochondria in cells, ranging from connected tubular structures to fragmented mitochondrial states [49, 98]. The proper distribution of mitochondria is essential for fulfilling the energy requirements of neurons [99]. As we age, our cells tend to accumulate more dysfunctional mitochondria that are unable to be removed due to diminishing mitophagy, hampered by the rise of oxidative stress [51]. Upon visualizing the mitochondrial morphology, we observed that all aged cell models had a more fragmented mitochondrial network morphology than young individuals. Additionally, due to the fragmented mitochondrial morphology, we observed an increase in the mitochondrial mass in the aged iNs and iPSCsNs. Knowing that the aged neurons reflected an impairment of mitochondrial properties, our findings indicated that an accumulation of damaged or aged mitochondria in both models of neuronal models was present in aging. Moreover, dysfunctional mitochondria tend to accumulate in the cells due to the decreased mitophagy mechanism [51, 100]. On the mRNA expression level, we observed an upregulation in the expression of the mitochondrial biogenesis-regulating gene, *NRF2*, in aged iNs, as well as in the aged iPSCsNs. Contrary to the belief that *NRF2* gene transcription declines with age [101–105], our findings in both neuronal models opposed this notion. Besides its role in mitochondrial biogenesis, NRF2 regulates the expression of genes for antioxidant, detoxifying, and cytoprotective proteins [106]. From this standpoint, the upregulation of NRF2 in both neuronal models could be considered as protective mechanism to counteract the rise of ROS. Our findings further presented an upregulation in fusion and fission regulating genes in the aged iNs but not in the aged iPSCsNs. Mitochondria can maintain their integrity by either promoting fission to remove damaged mitochondria or promoting fusion to exchange components, thereby reducing oxidative stress and strengthening the energy supply, which helps survival [107, 108]. Nevertheless, abnormalities in mitochondrial dynamics and quality control can lead to the accumulation of impaired and aged mitochondria, contributing to aging [108–110]. Given that the mitochondrial morphology impacts the bioenergetics and vice versa, a fragmented network morphology indicated a state of more damaged mitochondria and less mitochondrial metabolism, reflected in both aged neurons, iPSCsNs and iNs [48, 111].

On the mitochondrial level, we detected that aged iPSCsNs exhibit similar aging patterns to the aged iNs. Specifically, we spotted a decrease in bioenergetic properties, including impaired OXPHOS activity, which is linked to an overall reduction in cellular ATP levels. Additionally, the radical emission of mitochondrial superoxide anion and ROS reflected increased levels in both aged neuronal models. Furthermore, we observed an aging-associated imbalance in the redox state with a decrease in the redox ratio, accompanied by a drop in NAD^+^ and a rise in NADH. Regarding the mitochondrial network morphology, both aged neuronal models exhibited a fragmented mitochondrial network like the aged HFs.

It is worth noting that while aged iNs are known to preserve mitochondrial impairments, aged iPSCsNs are thought to lose their aging-associated donor signature [35, 39]. Our research, however, demonstrates that aged iPSCsNs can still preserve their donor-associated aging signature at the mitochondrial level, which is linked to their cell of origin. Furthermore, we exhibit a similar aging pattern as in the aged iNs at the mitochondrial level.

### 5.2. Reprogramming via the pluripotent states resets the metabolic shift towards anaerobic glycolysis linked to aging

Neurons are heavily dependent on the role of mitochondria as the powerhouse [44]. Nevertheless, as cells age, the functionality of these organelles diminishes, resulting in a deficiency of energy. In response to an energy shortage, the cells attempt to compensate for this deficit by switching to more anaerobic glycolytic metabolism [112]. However, this metabolic switch is unable to resolve the energy deficiency. Due to the smaller amount of ATP generated by glycolysis compared to mitochondrial OXPHOS (2 ATP versus 36 ATP), cells are unable to compensate for the lack of energy caused by mitochondrial function [97, 113–115]. The metabolic shift towards glycolysis and an increased glycolytic capacity in age might indicate that the aged cells adapted to the gradual manifestation of mitochondrial dysfunction by a higher glycolytic rate. In our cellular models, anaerobic glycolysis was more active in aged HFs and iNs than their younger counterparts. Our findings were consistent with other studies that observed a metabolism shift toward a glycolytic state due to mitochondrial impairments [112]. In this way, the cells were attempting to maintain a constant cellular ATP level in aging [42, 116, 117]. Contrary to the finding on the aged iNs, we did not exhibit an aging-associated metabolic switch towards anaerobic glycolysis in aged iPSCsNs by using the real-time assessment of anaerobic glycolysis. Upon comparing our findings to our previous study (unpublished data of the Eckert group) on aged iPSCs, we found that iPSCs from aged donors exhibit a metabolic shift towards anaerobic glycolysis compared to young donors. The production of lactate in the brain without the use of oxygen could be a way for the brain to survive when its ability to produce energy through the mitochondria is impaired [118]. As we age, the adult brain switches to anaerobic glycolysis to provide a limited energy source and time to maintain essential processes before the damage is too severe and death occurs. The LDHA enzyme tends to increase in older individuals, resulting in higher lactate levels, and is considered to be a marker of the anaerobic glycolytic pathway [119, 120]. In our investigation, it was shown that the levels of *LDHA* expression was significantly increased in aged iPSCsNs. The decline in mitochondrial function in aged iPSCsNs might cause a corresponding rise in the expression of the *LDHA* gene due to the energy shortage that occurred. Nevertheless, as these cells underwent various reprogramming stages (such as iPSCs and NPCs), their ability to adapt to the aging process and compensate for the energy deficit could be reset. Although the transcriptomic level of aged iPSCsNs was influenced by mitochondrial dysfunction, their glycolytic behavior did not demonstrate the same affection. Our speculation could be supported by our observation that there were no discernible differences in the mRNA expression of the *PFKFB3* gene between iPSCsNs of young and aged individuals. This suggests that the ability to transit to a more glycolytic metabolism was no longer present, perhaps due to a partial rejuvenation.

### 5.3. Strong discrepancies between aged iNs and aged iPSCsNs on the transcriptomic level

Previous studies have indicated that the induction of pluripotency resets age-related gene expression at the transcriptomic level, while direct conversion retains aging-associated donor signatures [23, 121]. As we observed mitochondrial impairments, we expected to observe similar transcriptomic profiles between aged iNs and aged iPSCsNs from the same donor. However, we presented that the population between the aged neuronal cell models differed strongly from each other. Moreover, we detected more significant differences in the aged iNs compared to young iNs. Our findings highlighted that impairments on functional properties after conversion were preserved, but the reprogramming via iPSCs-state induced a partial rejuvenation on the transcriptomic level. One example was the expression of the gene *PFKFB3*, which was upregulated in aged iNs compared to young iNs. However, the aged iPSCsNs showed no difference to iPSCsNs from young donors. PFKFB3 controls and sustains the levels of fructose-2,6-bisphosphate (F2,6P2), which is the most potent activator of 6-phosphofructo-1-kinase, a crucial regulator of glycolysis [122]. Neurons actively downregulate the expression of *PFKFB3* to maintain their antioxidant status by entering the PPP as an alternative route [44]. Neuronal aging is associated with an upregulation of *PFKFB3* expression, which enhances glycolysis (direct conversion of glucose to pyruvate) but also leads to cognitive impairments, synapse loss, and programmed cell death [123]. Other genes that showed significant regulation differences between the aged iNs and young iNs, as reflected in aging [54], but not in the aged iPSCsNs to the corresponding young were PARP1, AMPK, and AKT1 [47, 124]. Furthermore, the gene expression of *FOXO1* and *UPC2* exhibited an age-related gene expression level in aged iNs but not in aged iPSCsNs, where even the opposite gene regulation was spotted. Still, it has to be mentioned that there is controversy surrounding the aging-associated phenotype of UCP2, with some indicating an increase [125, 126] and others a decrease in regulation [127, 128]. UCP2 is an IMM protein vital in lowering MMP and dissipating metabolic energy to prevent oxidative stress [128]. The overall findings on the mRNA expression level suggest that differences between aged iNs and iPSCsNs from the same donors were present, indicating a vital link of rejuvenation on the mRNA level on the aged iPSCsNs.

### 5.4. Variations in Mitochondrial Properties and Aging Phenotype Among Individuals: Insights from a Comparative Analysis of Critical Parameters

The reasons behind this natural aging process have yet to be fully understood. Moreover, variations in the pace and intensity of aging among individuals exist [129]. When studying changes linked to the aging process of the human brain, it is essential to consider the concept of aging plasticity [6]. Curiously, the aging phenotype in the brain does not exhibit a uniform phenotype between individuals but instead shows significant variation from person to person [130]. Specific individuals demonstrate remarkable resilience as they age, while others exhibit a pronounced aging phenotype. The discrepancy between biological and chronological age must always be considered when conducting research [8]. This study conducted a comparative analysis of critical mitochondrial parameters to understand how the conversion process affects mitochondrial properties at the individual donor level. The results of this investigation revealed that the aging process was not uniform and showed, in some aspects, variation among the individuals and different cellular models. The findings indicated a notable correlation between distinct traits that persisted throughout the reprogramming process, and there were discernible discrepancies in certain traits among the individuals. Nevertheless, in direct and donor comparison, the aged iNs did not differ enormously from the iPSCsNs on the mitochondrial parameters. One exception in our investigation was the glycolytic capacity, as previously mentioned. Donor differences could be preserved within and between individuals after the reprogramming process [131]. One further distinction between the neuronal models and their corresponding HFs was that the aged HFs exhibited a more prominent ROS emission compared to the neuronal models. Upon seeing an increase in the mRNA level of NRF2 in aged neurons, a regulator of genes implicated in antioxidant processes [106], we hypothesize that some antioxidant mechanisms were activated in the aged neuronal models. Additionally, we observed increased gene expression of antioxidant proteins in both aged iPSCsNs and iNs, including *GPX1* and *CAT1*, respectively. Nevertheless, both processes of nuclear reprogramming increased the gene expression of antioxidant proteins. However, the underlying mechanism seemed unclear. In summary, these findings emphasized the significance of comprehending the adaptability of aging and the unique differences in how aging impacts mitochondrial function. In the investigation of aging, a direct comparison between two individuals could be insufficient to represent the physiological phenotype. Donor parameters might differ between donors, but most parameters showed a uniform phenotype among donors in the same age group. The results of this study might have consequences for the creation of focused interventions to decelerate or avert age-related mitochondrial dysfunction to utilize the aged iPSCsNs as a model system.

### 5.5. Telomerase activity in our neuronal models did not reflect the donor-associated decline in activity

The process of natural aging has traditionally been attributed to the initial factors of telomere shortening and mitochondrial dysfunction [8, 55]. While numerous theories elucidate the impact of various factors on the aging process, they often fail to account for the interconnection between these factors [132, 133]. Nevertheless, recent evidence suggested a significant connection between the shortening of telomeres and a decline in metabolic function [134]. The impaired functioning of mitochondria resulted in the gradual shortening of telomeres, which in turn triggered the reprogramming and malfunctioning of mitochondria [135]. This process has significant implications for the aging process and the development of various diseases. Gaining a comprehensive understanding of these interconnected mechanisms could potentially lead to the development of focused interventions aimed at reducing the effects of age-related diseases. Telomerase, the enzyme accountable for preserving the length of telomeres, has been discovered to possess a protective function in the mitochondria, specifically in situations involving oxidative stress [136]. There is growing evidence that telomerase can aid cell survival and stress resistance in a way that is not dependent on telomeres. Telomerase is known to move around within cells. When oxidative stress levels increase, telomerase can be detected in the mitochondria. This trait is associated with lower oxidative stress and improved mitochondrial function in cells expressing telomerase, although the mechanisms behind this are not fully understood. Our observation revealed the presence of telomere attrition, a well-recognized modification linked with aging, in aged HFs. This was shown by a reduction in telomere length and a decrease in telomerase activity compared to young individuals. Nevertheless, we have seen no disparity in the telomerase activity between the aged neuronal cell models, iNs and iPSCsNs, and their young counterparts. In general, we speculate that this might be due to the fact that neurons do not display a pronounced TA. It is commonly believed that TA is limited to dividing cells, whereas non-dividing cells, such as neurons, may not require TA anymore. However, research has shown that telomerase proteins may also be produced in post-mitotic neurons, albeit at a lower rate than actively dividing cells [59, 137, 138]. To detect a sufficient rate of telomerase activity, we needed to increase the number of neurons to a cell count of 100’000, whereas, for HFs and iPSCs, a cell count of 10’000 was necessary. The suitability of our TRAP assay combined with qPCR for identifying telomerase activity in neurons remains uncertain, given that the low telomerase activity in neurons might not enable substantial changes to be quantified [139]. Our research has further revealed some noteworthy findings regarding the TL and TA in young and aged iPSCs. Notably, we observed no significant differences in the telomere length and telomerase activity between the young and aged iPSCs. However, we observed a prolongation of the TL after the conversion mechanism to the iPSCs state. This observation aligns with prior research showing that telomeres in iPSCs are longer than those in their parental cell lines [29, 140, 141].

## 6. Conclusion

In summary, our findings indicate that aged iNs and aged iPSCsNs from the *same aged cells of origin*, HFs, exhibit a wide array of aging phenotypes linked to mitochondria that have been previously observed in *in vivo* aging. For model human neuronal aging, iNs are a promising model system. Nevertheless, technical limitations exist when using iNs, especially when considering the cell number of neurons at the end [64, 142]. Hereby, we could demonstrate that the aged iPSCsNs could be a good replacement model in maintaining an aging-associated impairment on the mitochondrial level. Although our use of aged iPSCsNs and iNs represented a significant step forward in modeling neuronal aging *in vitro,* it is yet to be determined whether these methods accurately replicate natural human neuronal aging. Nonetheless, we were able to demonstrate an aging phenotype in the aged iPSCsNs similar to the aged iNs on mitochondrial phenotypes. However, regarding mRNA expression level and anaerobic glycolysis, the aged iPSCsNs could not reflect a phenotype like that observed in the aged iNs. This difference could be attributed to the different nuclear reprogramming approaches and might indicate that the aged iPSCsNs could experience a partial rejuvenation on some parameters. However, there is a need for more research examining the maintenance of the aging characteristics in iPSCsNs and iNs. While the majority of research has focused on age-related neurodegenerative diseases, expanding our knowledge of the underlying disease-specific molecular pathways, significantly fewer studies targeted molecular alteration of the aging brain in the absence of diseases. In the pathological field, conceptually similar work has been carried out by Jiao and the group comparing iPSCsNs and aged iNs from Dravet syndrome patients (severe myoclonic epilepsy of infancy) [143]. Their findings presented that both neuronal models could represent a hyperexcitable state linked to the pathological phenotype. Thereby presenting that the iPSCsNs could recapitulate the neuronal pathology. Further publication around iPSCsNs derived from patients with Alzheimer’s disease exhibited high levels of pathological markers linked to Alzheimer’s disease [144–146]. Still, it was highlighted that prolonged cultivation with astrocytes might be required to study Alzheimer’s disease-associated alteration, including loss of synaptic proteins. These attempts using iPSCsNs to investigate pathological phenotypes have offered an essential demonstration of the effectiveness of these cells in replicating the underlying biochemical mechanism. Regardless, research demonstrating a preservation of the aging or pathological phenotype in the iPSCs or iPSCsNs is rare. In summary, our work enhanced our comprehension of the extent to which iNs or iPSCsNs derived from aged donors accurately simulate the process of human brain aging by *in vitro* modeling. The aged iPSCsNs exhibit an aging phenotype like aged iNs, making them a valuable resource for studying bioenergetics and performing drug screening. These applications need a substantial quantity of material, making the aged iPSCsNs a viable tool due to their possession of aging-associated markers. However, aged iNs are more favorable *in vitro* neuronal models by preserving an overall aging signature of the donor beyond mitochondrial dysfunction. Still, further research is required to understand the mechanism responsible for maintaining aging characteristics.

## 7. Material and Methods

### 7.1. Reagents and Tools

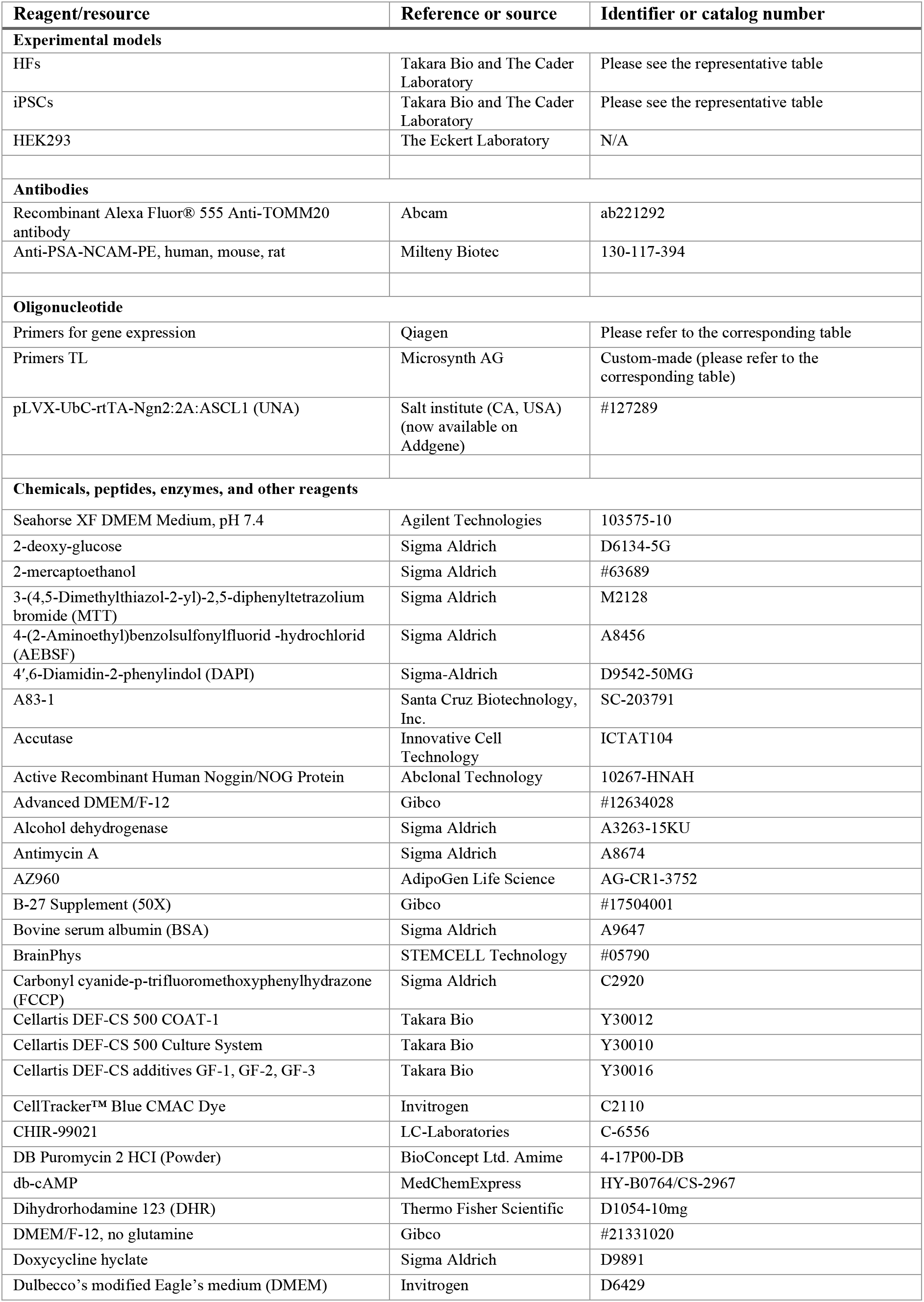
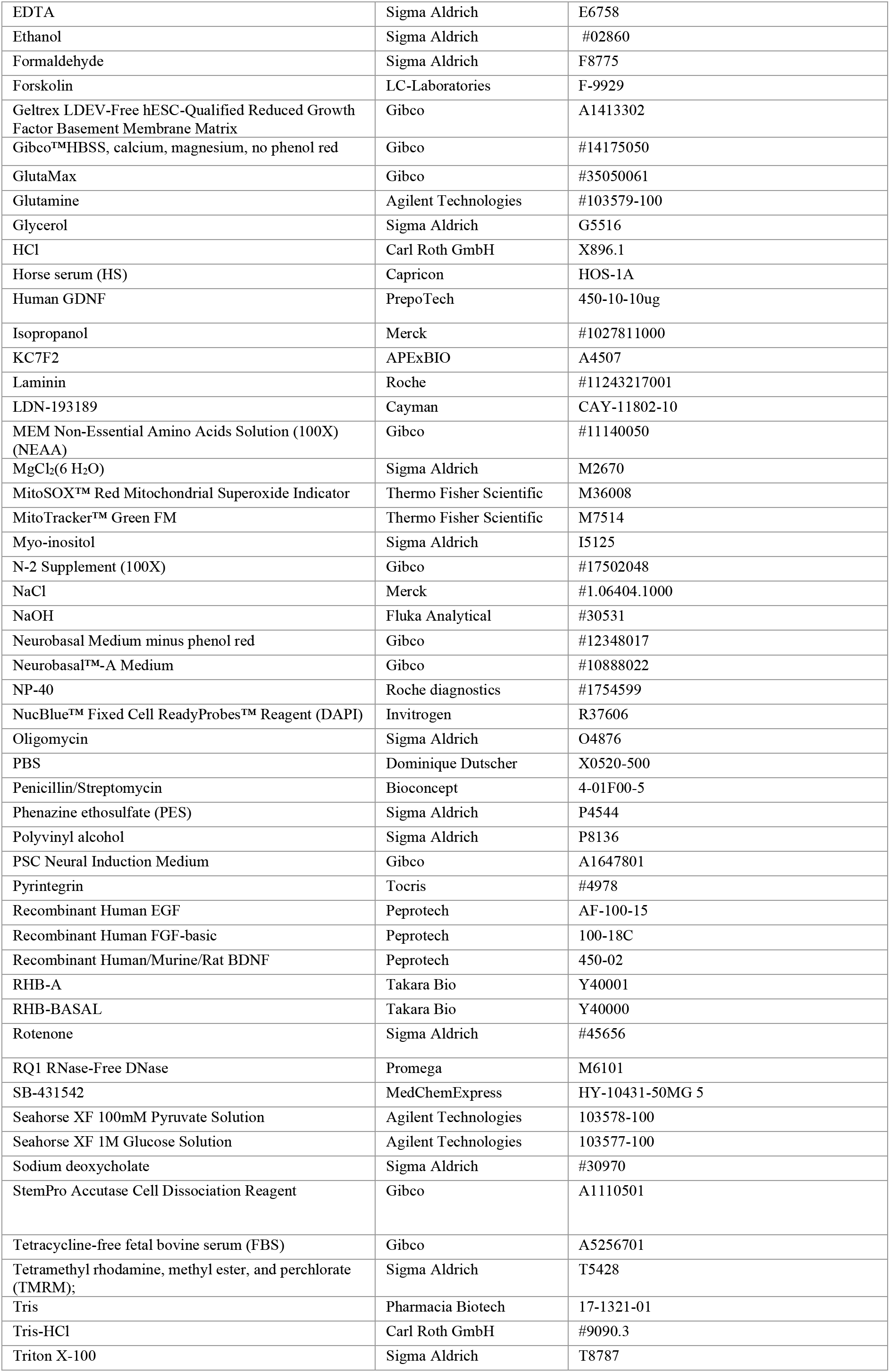
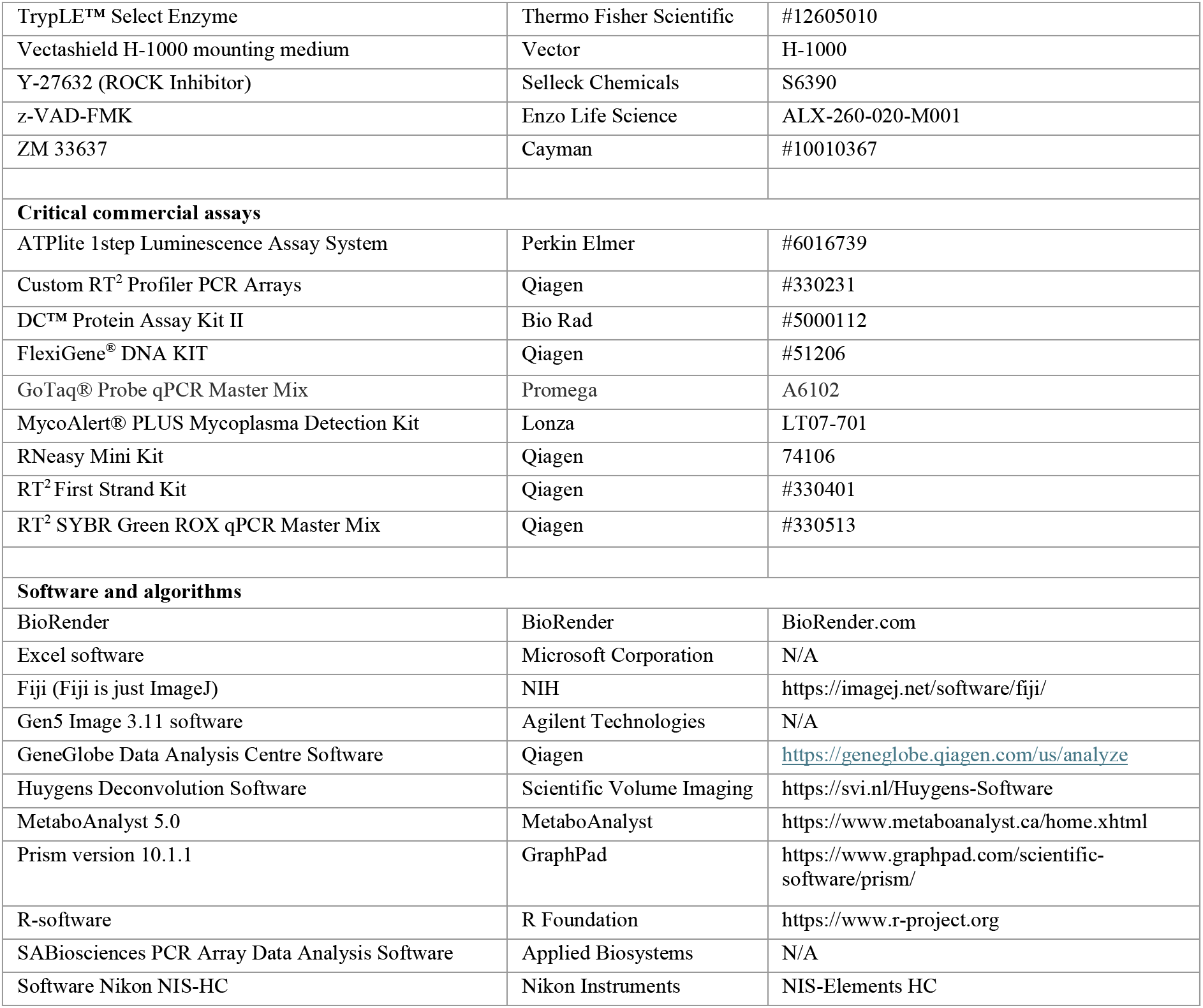

| Reagent/resource | Reference or source | Identifier or catalog number |
| --- | --- | --- |
| **Experimental models** |  |  |
| HF<sub>s</sub> | Takara Bio and The Cader Laboratory | Please see the representative table |
| iPSCs | Takara Bio and The Cader Laboratory | Please see the representative table |
| HEK293 | The Eckert Laboratory | N/A |
| **Antibodies** |  |  |
| Recombinant Alexa Fluor® 555 Anti-TOMM20 antibody | Abcam | ab221292 |
| Anti-PSA-NCAM-PE, human, mouse, rat | Milteny Biotec | 130-117-394 |
| **Oligonucleotide** |  |  |
| Primers for gene expression | Qiagen | Please refer to the corresponding table |
| Primers TL | Microsynth AG | Custom-made (please refer to the corresponding table) |
| pLVX-UbC-rtTA-Ngn2:2A:ASCL1 (UNA) | Salt institute (CA, USA) (now available on Addgene) | #127289 |
| **Chemicals, peptides, enzymes, and other reagents** |  |  |
| Seahorse XF DMEM Medium, pH 7.4 | Agilent Technologies | 103575-10 |
| 2-deoxy-glucose | Sigma Aldrich | D6134-5G |
| 2-mercaptoethanol | Sigma Aldrich | #63689 |
| 3-(4,5-Dimethylthiazol-2-yl)-2,5-diphenyltetrazolium bromide (MTT) | Sigma Aldrich | M2128 |
| 4-(2-Aminoethyl)benzolsulfonylfluorid -hydrochlorid (AEBSF) | Sigma Aldrich | A8456 |
| 4',6-Diamidin-2-phenylindol (DAPI) | Sigma-Aldrich | D9542-50MG |
| A83-1 | Santa Cruz Biotechnology, Inc. | SC-203791 |
| Accutase | Innovative Cell Technology | ICTAT104 |
| Active Recombinant Human Noggin/NOG Protein | Abclonal Technology | 10267-HNAH |
| Advanced DMEM/F-12 | Gibco | #12634028 |
| Alcohol dehydrogenase | Sigma Aldrich | A3263-15KU |
| Antimycin A | Sigma Aldrich | A8674 |
| AZ960 | AdipoGen Life Science | AG-CR1-3752 |
| B-27 Supplement (50X) | Gibco | #17504001 |
| Bovine serum albumin (BSA) | Sigma Aldrich | A9647 |
| BrainPhys | STEMCELL Technology | #05790 |
| Carbonyl cyanide-p-trifluoromethoxyphenylhydrazone (FCCP) | Sigma Aldrich | C2920 |
| Cellartis DEF-CS 500 COAT-1 | Takara Bio | Y30012 |
| Cellartis DEF-CS 500 Culture System | Takara Bio | Y30010 |
| Cellartis DEF-CS additives GF-1, GF-2, GF-3 | Takara Bio | Y30016 |
| CellTracker™ Blue CMAC Dye | Invitrogen | C2110 |
| CHIR-99021 | LC-Laboratories | C-6556 |
| DB Puromycin 2 HCl (Powder) | BioConcept Ltd. Amime | 4-17P00-DB |
| db-cAMP | MedChemExpress | HY-B0764/CS-2967 |
| Dihydrorhodamine 123 (DHR) | Thermo Fisher Scientific | D1054-10mg |
| DMEM/F-12, no glutamine | Gibco | #21331020 |
| Doxycycline hyclate | Sigma Aldrich | D9891 |
| Dulbecco's modified Eagle's medium (DMEM) | Invitrogen | D6429 |
| EDTA | Sigma Aldrich | E6758 |
| Ethanol | Sigma Aldrich | #02860 |
| Formaldehyde | Sigma Aldrich | F8775 |
| Forskolin | LC-Laboratories | F-9929 |
| Geltrex LDEV-Free hESC-Qualified Reduced Growth Factor Basement Membrane Matrix | Gibco | A1413302 |
| Gibco™HBSS, calcium, magnesium, no phenol red | Gibco | #14175050 |
| GlutaMax | Gibco | #35050061 |
| Glutamine | Agilent Technologies | #103579-100 |
| Glycerol | Sigma Aldrich | G5516 |
| HCl | Carl Roth GmbH | X896.1 |
| Horse serum (HS) | Capricorn | HOS-1A |
| Human GDNF | PreproTech | 450-10-10ug |
| Isopropanol | Merck | #1027811000 |
| KC7F2 | APExBIO | A4507 |
| Laminin | Roche | #11243217001 |
| LDN-193189 | Cayman | CAY-11802-10 |
| MEM Non-Essential Amino Acids Solution (100X) (NEAA) | Gibco | #11140050 |
| MgCl<sub>2</sub>(6 H<sub>2</sub>O) | Sigma Aldrich | M2670 |
| MitoSOX™ Red Mitochondrial Superoxide Indicator | Thermo Fisher Scientific | M36008 |
| MitoTracker™ Green FM | Thermo Fisher Scientific | M7514 |
| Myo-inositol | Sigma Aldrich | I5125 |
| N-2 Supplement (100X) | Gibco | #17502048 |
| NaCl | Merck | #1.06404.1000 |
| NaOH | Fluka Analytical | #30531 |
| Neurobasal Medium minus phenol red | Gibco | #12348017 |
| Neurobasal™-A Medium | Gibco | #10888022 |
| NP-40 | Roche diagnostics | #1754599 |
| NucBlue™ Fixed Cell ReadyProbes™ Reagent (DAPI) | Invitrogen | R37606 |
| Oligomycin | Sigma Aldrich | O4876 |
| PBS | Dominique Dutscher | X0520-500 |
| Penicillin/Streptomycin | Bioconcept | 4-01F00-5 |
| Phenazine ethosulfate (PES) | Sigma Aldrich | P4544 |
| Polyvinyl alcohol | Sigma Aldrich | P8136 |
| PSC Neural Induction Medium | Gibco | A1647801 |
| Pyrintegrin | Tocris | #4978 |
| Recombinant Human EGF | Peprotech | AF-100-15 |
| Recombinant Human FGF-basic | Peprotech | 100-18C |
| Recombinant Human/Murine/Rat BDNF | Peprotech | 450-02 |
| RHB-A | Takara Bio | Y40001 |
| RHB-BASAL | Takara Bio | Y40000 |
| Rotenone | Sigma Aldrich | #45656 |
| RQ1 RNase-Free DNase | Promega | M6101 |
| SB-431542 | MedChemExpress | HY-10431-50MG 5 |
| Seahorse XF 100mM Pyruvate Solution | Agilent Technologies | 103578-100 |
| Seahorse XF 1M Glucose Solution | Agilent Technologies | 103577-100 |
| Sodium deoxycholate | Sigma Aldrich | #30970 |
| StemPro Accutase Cell Dissociation Reagent | Gibco | A1110501 |
| Tetracycline-free fetal bovine serum (FBS) | Gibco | A5256701 |
| Tetramethyl rhodamine, methyl ester, and perchlorate (TMRM); | Sigma Aldrich | T5428 |
| Tris | Pharmacia Biotech | 17-1321-01 |
| Tris-HCl | Carl Roth GmbH | #9090.3 |
| Triton X-100 | Sigma Aldrich | T8787 |
| TrypLE™ Select Enzyme | Thermo Fisher Scientific | #12605010 |
| Vectashield H-1000 mounting medium | Vector | H-1000 |
| Y-27632 (ROCK Inhibitor) | Selleck Chemicals | S6390 |
| z-VAD-FMK | Enzo Life Science | ALX-260-020-M001 |
| ZM 33637 | Cayman | #10010367 |
| **Critical commercial assays** |  |  |
| ATPlite 1step Luminescence Assay System | Perkin Elmer | #6016739 |
| Custom RT<sup>2</sup> Profiler PCR Arrays | Qiagen | #330231 |
| DC™ Protein Assay Kit II | Bio Rad | #5000112 |
| FlexiGene® DNA KIT | Qiagen | #51206 |
| GoTaq® Probe qPCR Master Mix | Promega | A6102 |
| MycoAlert® PLUS Mycoplasma Detection Kit | Lonza | LT07-701 |
| RNeasy Mini Kit | Qiagen | 74106 |
| RT<sup>2</sup> First Strand Kit | Qiagen | #330401 |
| RT<sup>2</sup> SYBR Green ROX qPCR Master Mix | Qiagen | #330513 |
| **Software and algorithms** |  |  |
| BioRender | BioRender | BioRender.com |
| Excel software | Microsoft Corporation | N/A |
| Fiji (Fiji is just ImageJ) | NIH | https://imagej.net/software/fiji/ |
| Gen5 Image 3.11 software | Agilent Technologies | N/A |
| GeneGlobe Data Analysis Centre Software | Qiagen | https://geneglobe.qiagen.com/us/analyze |
| Huygens Deconvolution Software | Scientific Volume Imaging | https://svi.nl/Huygens-Software |
| MetaboAnalyst 5.0 | MetaboAnalyst | https://www.metaboanalyst.ca/home.xhtml |
| Prism version 10.1.1 | GraphPad | https://www.graphpad.com/scientific-software/prism/ |
| R-software | R Foundation | https://www.r-project.org |
| SABiosciences PCR Array Data Analysis Software | Applied Biosystems | N/A |
| Software Nikon NIS-HC | Nikon Instruments | NIS-Elements HC |

### 7.2. Cell culture

The primary human fibroblasts (HFs) and the corresponding iPSCs were purchased from Takara Bio (Kusatsu, Shiga, Japan) or were kindly provided by Dr. Zameel Cader (University of Oxford) and the Stem cells for biological assays of novel drugs and predictive toxicology (StemBANCC) consortium. For an in-depth comprehension of donor information, please see Table 1.

The iPSCsNs and iNs were generated at the Neurobiology Lab for Brain Aging and Mental Health (Basel). All cells were constantly checked for mycoplasma using the MycoAlert® PLUS Mycoplasma Detection Kit.

#### 7.2.1. Primary Human Fibroblasts (HFs)

The HFs were cultured in a growing medium (GM) composed of DMEM supplemented with 1 % Penicillin-Streptomycin, 1 % Glutamax, and 20 % tetracycline-free FBS. The cultivation process was carried out at 37 °C and 5 % CO_2_ in a humidified incubator. The HFs were grown on 10 cm^2^ dishes and were split at a ratio of 1:3 at a confluency of 80 % -100 %. Since HFs have a circadian rhythm [111], it was essential to synchronize the HFs one day before carrying out the experiments. The synchronization was accomplished using a serum shock technique, in which the HFs were exposed to GM supplemented with 50 % horse serum for a period of 2 hours at 37°C. HFs were platted into either FBS pre-coated 96-well plates at a density of 1.5 x 10^4^ cells per well, FBS pre-coated XF24 cell culture microplate plates at a density of 2 x 10^4^ cells per well, or 12-well plates with coverslips at a density of 1 x 10^4^ cells per well, depending on the experiments.

#### 7.2.2. Directly converted neurons (iNs) from primary human fibroblasts

The iNs were obtained from HFs following the approach previously outlined by Zhou-Yang and colleagues, with some minor adjustments [147]. To generate iNs from HFs, the HFs were transduced with a puromycin-resistant lentiviral vector that carried the ‘all-in-one’ vector pLVX-UbC-rtTA-Ngn2:2A:ASCL1 (UNA), which also compose a resistance gene to puromycin. The transduction was performed in TMF medium composed of DMEM with 20 % tetracycline-free fetal bovine serum and 0.1 % NEAA. For the expansion, the transduced HFs were cultured in TMF containing puromycin (1 mg/mL) for at least five passages after viral transduction. For the neuronal conversion, 100 % confluent UNA-transduced HFs were pooled into higher density by 3:1 pooling split on 6-well plates. One day after the pooling, the medium was exchanged to neuronal conversion medium (NK-medium) containing DMEM/F-12, Neurobasal-A, 1xB-27, 1xN-2, 1 μg/ml Laminin, 400 μg/ml db-cAMP, 2 μg/ml doxycycline, 150 ng/ml Noggin, 0.5 μM LDN-193189, 0.5 μM A83-1, 3 μM CHIR-99021, 5 μM Forskolin, 10 μM SB-431542, 1 μM Pyrintegrin, 7,5 μM KC7F2, 0.1 μM AZ960, and 0,75 μM ZM33637. The NK medium was replaced every other day for a duration of 3-4 weeks. Following the conversion, the cells were dissociated with TrypLE™ Select Enzyme and sorted using FACS. The iNs were sorted by positive PSA-NCAM-PE and negative DAPI staining in sorting buffer consisting of 150 mM myo-inositol, 5 mg/mL polyvinyl alcohol, 1 % DNase, and 10 μM Rock-inhibitor in PBS by using the FACS Sorter Aria III (FACS Core Facility, Biozentrum, University of Basel). The sorted iNs were gathered in NK-medium containing Rock-inhibitor (Y-27632) and z-VAD-FMK and plated either on Geltrex precoated μ-Slide 8 well ibidi chambers, Seahorse XFp Cell Culture Miniplate at a density of 2,5 x 10^4^ cell/well or on 96-well plates at a density of 5 x 10^4^ cell/well depending on the experiments. The next day, the medium was replaced with neuronal maturation media consisting of BrainPhys, 1xB-27, 1xN-2, 1 μg/ml Laminin, 500 μg/ml db-cAMP, 20 ng/ml GDNF, and 20 ng/ml BDNF and refreshed every 24 hours. Following 72 hours of plating, the neural connections were regenerated, and measurements were performed.

#### 7.2.1. Induced pluripotent stem cells (iPSCs)

The iPSCs were cultured in feeder-free condition and on Cellartis DEF-CS COAT-1-coated plates using the Cellartis DEF-CS culture system, following the manufacturer’s instructions of Takara Bio and previously described [148]. The iPSCs were maintained in a 37°C, 5 % CO_2_ humidified incubator with daily medium replacement. Passaging was done twice a week using TripLE Select.

#### 7.2.2. Induced pluripotent stem cell derived neurons (iPSCsNs)

Based on the instructions provided by the manufacturer Gibco using the PSC Neural Induction protocol, the neural induction process of young and aged iPSCs to neural progenitor cells (NPCs) was conducted. Initially, iPSCs were cultured onto Cellartis DEF-CS COAT-1-coated 6-well cell culture plates at a density of 2.6 x 10^5^ cells per well using the Cellartis DEF-CS culture system. At the 6-hour mark (day 0), the medium was replaced with neural induction medium (NIM), consisting of Neurobasal Medium minus phenol red and 2 % Neural Induction Supplement. The NIM was replaced every other day from day 0 to day 4 of neural induction and changed daily after day 4 when the cells attained confluency. On the seventh day, primitive NPCs were dissociated using StemPro Accutase and placed onto Geltrex-coated 6-well cell culture plates. The cells were added at a density of 4.8-9.6 x 10^5^ cells per well in neural expansion medium (NEM). The NEM consisted of a 1:1 ratio of Neurobasal Medium to Advanced DMEM/F-12 with 2 % Neural Induction Supplement and 10 μM Y-27632. The NEM was replaced every two days until the NPCs acquired confluency. In the following step, the expanded NPCs were either cryopreserved in NEM with 10 % DMSO or converted into iPSCsNs using the protocol of Takara Bio. To stimulate the conversion from NPCs to iPSCsNs, the NPCs were cultivated on Geltex-coated 6-well plates using RHB-A media, which was enhanced with 20 ng/mL EGF and 20 ng/mL FGF for a minimum of two passages. StemPro Accutase was used for passaging NPCs (not more than five passages) and additionally treated overnight with 10 μM ROCK Inhibitor after splitting. To convert NPCs into iPSCsNs, StemPro Accutase was first used to dissociate NPCs, which were then placed in Geltrex-coated 10 cm^2^ cell culture dishes with a density of 4.5 x 10^5^ cells per dish. The cells were cultured in RHB-BASAL medium supplemented with 0.5 % NDiff N2, 1 % B-27 Supplement, and 10 ng/mL FGF. Additionally, for one night, 10 μM Y-27632 was added. Every other day for a duration of 6 days, half of the culture medium was substituted with fresh medium. On the seventh day, the differentiation medium was changed to a mixture of RHB-BASAL medium and Neurobasal Medium without phenol red in a 1:1 ratio. This mixture was supplemented with 0.25 % 1xN-2, 1 % B-27 Supplement, 10 ng/mL FGF, and 0.5 % GlutaMAX. Every other day until day 13, half of the cultural media was refreshed. On the 14th day, the differentiation medium was changed to Neurobasal Medium without phenol red, adding 2 % B-27 Supplement, 1 % GlutaMAX, and 20 ng/mL BDNF. The cells were incubated for 14 days, with half of the culture media replaced every other day. The iPSCsNs were transferred to the appropriate assay plates to conduct various investigations. In summary, iPSCsNs were separated using StemPro Accutase with 10 μM Y-27632 for 45 minutes at 37°C. They were then transferred to either Geltrex-coated 96-well cell culture plates at a density of 6.0 x 10^5^ cells per well, Geltrex-coated 96-well plates at a density of 6.0 x 10^5^ cells per well, or Geltrex-coated Seahorse XFp Cell Culture Miniplate at 3.0 x 10^5^ cells per well. The iPSCsNs were cultured in Neurobasal Medium without phenol red, supplemented with 2 % B-27 Supplement, 1 % GlutaMAX, and 20 ng/mL BDNF with 10 μM of the ROCK Inhibitor only for an overnight addition. Every other day for seven days before conducting the experiments, half of the culture media was replaced with fresh medium.

### 7.3. Cellular ATP level

For the determination of the total ATP content in the cells, the bioluminescence assay ATPlite 1step was conducted following the manufacturer’s instruction [149]. The bioluminescence method measures the light produced by the luciferase reaction of ATP and luciferin, presenting proportional to the ATP concentration of the cells. The cells were plated on pre-coated transparent 96-well cell culture plates at the corresponding density. The emitted light was measured using the Cytation 3 Cell Imaging Multi-Mode Reader.

### 7.4. Mitochondrial Membrane Potential (MMP)

The fluorescent dye TMRM was used to measure the MMP [148]. The cells were seeded at the appropriate densities onto black 96-well cell culture plates and incubated with 0.4 μM TMRM for 30 min at 37 °C and 5 % CO_2_ in the incubator. Subsequently, the cells underwent two rounds of washing with HBSS. The fluorescence at 548 nm (excitation) and 574 nm (emission) was then measured using the Cytation 3 Cell Imaging Multi-Mode Reader. The spatial arrangement of the dye across the cell membrane directly indicates the level of MMP.

### 7.5. Superoxide Anion Radical Levels and Mitochondrial Reactive Oxygen Species (ROS)

Different fluorescent dyes were used to determine ROS production and superoxide anion radicals. Specifically, DHR was used to measure the mitochondrial ROS level, while MitoSOX was used to detect the mitochondrial superoxide anion radical level (°O_2_^−^)) [111]. The cells were carefully plated onto black 96-well cell culture plates and then incubated with 10 μM of DHR for 30 minutes or 5 μM of MitoSOX for 2 hours at 37 °C and a 5 % CO_2_ in the incubator. Following incubation, the cells were rinsed twice with HBSS before measurement. The Cytation 3 Cell Imaging Multi-Mode Reader was then used to measure DHR at 485 nm (excitation)/535 nm (emission) and MitoSOX at 531nm (excitation)/595 nm (emission).

### 7.6. NAD^+^ to NADH

To measure the intracellular NAD^+^/NADH content, an enzyme cycling assay was performed [111, 150, 151]. The cells were lysed in 250 μl of protein lysis buffer (150 mM Tris, 150 mM NaCl, 1 % NP-40, 0.1 % SDS, and 2 mM EDTA) for the sample preparation. In the next step for the NAD^+^ extraction, 100 μl lysate was heat-incubated with 100 μl HCl (0.1 M) for 5 min at 95°C and shaken at 300 rpm, followed by cooling the sample on ice. In the next step, the NAD^+^ extract was neutralized by adding 100 μl of NaOH (1 M) and then centrifuged at 10’000 g for 10 min at 4 °C. The supernatant was collected and placed on ice or was stored at –80°C for a later continuation. For the NADH extraction, the same step as for the NAD^+^ extraction was carried out with the difference in the heat-incubation with HCl; the sample was heat-incubated with NaOH (1 M) and then neutralized with HCl (1M). With the remaining lysate, the protein content was determined (see protein assay). For the cycling assay, 50 μl of the samples or the corresponding standards, either NAD^+^ or NADH (range from 20 nM to 400 nM), were plated on a transparent 96-well plate. The samples were then incubated with 50 μl of a mixture containing: 0,02 M Tricine-NaOH buffer (pH 8), 8 mM EDTA, 0.84 mM MTT, 3,32 mM PES, and 1 M Ethanol for 10 min at 37°C. After this step, alcohol dehydrogenase (10 U/ml) was added and incubated at 37°C for 1 hour. Following this, the reaction was stopped by adding 50 μl NaCl (6 M). In the last step, 100 μl ethanol (96 %) was mixed in every well to solubilize the formazan. The principle behind this enzymatic cycling assay is that the electron in the acid-base extracted sample is transferred from ethanol over reduced pyridine nucleotides to an MTT coupled reaction and lastly to PES, then to be visualized by the purple formazan formation. To detect the crystal formation, the absorbance was detected at 595 nm using the Cytation 3 Cell Imaging Multi-Mode Reader.

#### 7.6.1. Determination of Protein content – Lowry assay

The protein content was determined using the DCTM Protein Assay Kit and proceeded following the manufacturing script [148]. Therefore, the lysate samples were diluted 1 to 10 in PBS^+ Mg, + Ca^ and a standard of BSA in the range of 2 mg/ml to 0.03125 mg/ml was prepared. Then, 5 μl of diluted sample and standard was loaded on the respective well on a 96-well plate. Following this step, 25 μl of reaction A and 200 μl of reaction B were added to each well. At room temperature, the plate was shaken at 180 rpm for 10 minutes. The absorbance was then read at 690 nm using the Cytation 3 Cell Imaging Multi-Mode Reader.

### 7.7. Mitochondrial Respiration and Glycolysis

The Seahorse analyzers were used to measure critical parameters of mitochondrial respiration and anaerobic glycolysis. The Seahorse analyzer enables real-time quantification of oxygen consumption rate (OCR) directly linked to mitochondrial respiration. It also measures the extracellular acidification rate (ECAR), indicating the anaerobic glycolysis activity level. In this context, the sequential injection of mitochondrial modulators enabled the examination of the following key elements of mitochondrial respiration properties: basal respiration, ATP turnover, maximal respiration, and respiratory reserve capacity (Table 2) [111, 152]. The sequential administration of glycolytic modulators enabled the identification of the key parameters of glycolysis, including glycolysis itself, glycolytic capacity, and non-glycolytic acidification (Table 3).

**Table 2:**
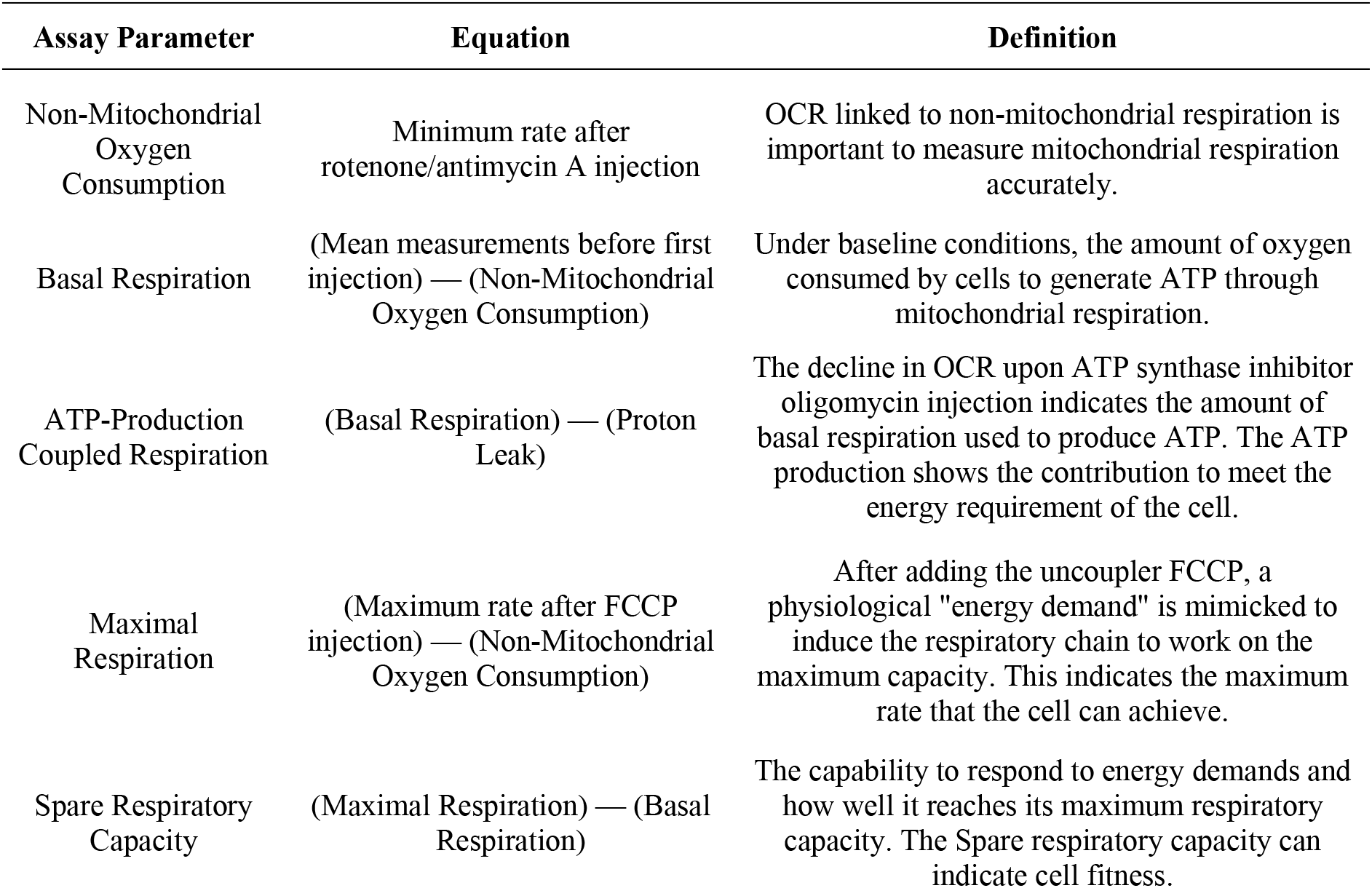
Mitochondrial stress test parameters. FCCP: carbonyl cyanide-p-trifluoromethoxyphenylhydrazone; OCR: oxygen consumption rate.

**Table 3:** Glycolysis stress test parameters. ECAR: extracellular acidification rate; OXPHOS: oxidative phosphorylation; 2-DG: 2-deoxy-glucose.

| Assay Parameter | Equation | Definition |
| --- | --- | --- |
| Non-Glycolytic Acidification | Minimum rate measurement after 2-DG injection | This indicated the ECAR not linked to the glycolysis. Essential to determine accurately the glycolysis parameter. |
| Basal glycolysis or glycolysis | (Maximum rate measurement between glucose and oligomycin injection) — (Non-Glycolytic Acidification) | The process of measuring glycolysis by converting glucose to pyruvate after adding a saturating amount of glucose. |
| Glycolytic Capacity | (Maximum rate measurement between Oligomycin and 2-DG injection) — (Non-Glycolytic Acidification) | The maximal glycolysis capacity by shutting down the OXPHOS and forcing the cell to use glycolysis at its maximal capacity. |

The Seahorse XFe24 Analyzer was used to measure the HFs, whilst the Seahorse XF HS Mini Analyzer was employed for both neuronal models. Before taking measurements, the medium was replaced with the assay medium, which consisted of Seahorse XF DMEM medium, pH 7.4. For measuring the OCR, the medium was supplemented with 25 mM glucose, 4 mM glutamine, and 1 mM pyruvate for HFs and iNs, whereby for iPSCsNs a concentration of 18 mM glucose, 4 mM glutamine, and 2 mM glutamine were added. To measure the ECAR, the glucose was not added in the assay medium. Before taking the measurements, the seahorse plates were equilibrated in a CO_2_-free incubator at 37°C for 45-60 minutes. The XF Mito Stress Test methodology was executed in accordance with the manufacturer’s instructions to evaluate the essential parameters of mitochondrial respiration. The OCR rate was recorded under basal condition followed by sequential injection of 1 μM (HFs), 2.5 μM (iNs), or 1,5 μM (iPSCsNs) oligomycin, 2 μM (HFs & iNs) or 1 μM (iPSCsNs) FCCP, and a combination of 4 μM (HFs), 2 μM (iNs), or 0,5 μM antimycin A and 2 μM (HFs & iNs) or 1 μM (iPSCsNs) rotenone. The Seahorse XF Glycolysis Stress Test was conducted to identify glycolysis by detecting the ECAR, following the directions provided by the manufacturer. The ECAR was measured under basal condition, followed by 25 mM glucose (HFs & iNs) or 18 mM (iPSCsNs), 1 μM (HFs), 2.5 μM (iNs), or 1,5 μM (iPSCsNs) oligomycin, and 25 mM (HFs & iNs) or 50 mM 2-deoxy-glucose by sequential injection. The critical parameters were computed manually, and the calculations are documented in table 2 for the Mitochondrial Stress Test and in table 3 for the Glycolysis Stress Test.

### 7.8. Mitochondrial network morphology

To observe the structure of the mitochondrial network morphology, HFs were plated onto FBS-precoated coverslips and both neuronal models onto geltrex precoated μ-Slide 8 well ibidi chambers or 96-well plates. The assessment was conducted as previously described [148]. The different cell types were fixed with 4 % formaldehyde for 15 min and then rinsed twice with PBS^+ Mg, + Ca^. To permeabilize the cells 0.2 % triton X-100 for 15 minutes was used and then blocked with 2 % BSA for 1 hour. Then, the mitochondria were stained with the mitochondrial marker translocase of the outer mitochondrial membrane complex subunit 20, TOMM20 Alexa Fluor^®^ 555, for 4 hours at room temperature. The coverslip with the HFs were mounted using the Vectashield H-1000 mounting medium and the neurons were visualized in PBS. The microscopy images for HFs and iNs were obtained using an inverted microscope (Leica Microsytemes TCS SPE DMI4000) connected to an external light source (Leica EL6000). The images were taken using an x63 oil immersion objective. Before examining the mitochondrial structure in the iNs, the images were subjected to deconvolution using Huygens Deconvolution Software to minimize non-specific signals. The iPSCsNs were visualized by the Nikon Inverted Research Microscope Eclipse Ti2-E using the x60 oil immersion objective. Further, the iPSCsNs were stained with NucBlue™ Fixed Cell ReadyProbes™ Reagent (DAPI) to visualize the nucleus, and the Nikon NIS-HC software was used. An acquisition of XYZ-confocal image with maximum intensity projection was utilized for all cell types. The acquisition settings within each cell type were consistent throughout the imaging process. The morphological investigation included blind examination using the NIH ImageJ program [153], which was applied to the whole image. The changes on the analysis settings were set for the rolling ball as 5 for iNs and HFs, whereas for iPSCsNs, a rolling ball 10 was chosen. The criteria used to describe the morphology of mitochondria include Form Factor (FF), Area-Weighted Form Factor (AW), Aspect Ratio (AR), and mitochondrial length. The FF, also known as mitochondrial elongation, quantifies the intricate form of the particle by measuring its circularity in an inverted manner. The AW is a variant of the FF that exhibits more reliability when dealing with mitochondria closely positioned near each other. In contrast, the FF is more dependable when dealing with wide-spaced mitochondria. The AR indicated the major-to-minor axis ratio without considering the perimeter. A value of 1 is regarded as a perfect circle for the FF, AW, and AR. The Mitochondrial length indicates the particle length in units of pixels after the mitochondria are minimized to a single-pixel-wide shape. Hence, the area, perimeter, major axis, and minor axis measurements are disregarded. The combination of visual inspection and categorical scoring is important for assessing changes in mitochondrial network morphology. In general, all data were obtained within the dynamic range of each camera, and all measurements were conducted on the unprocessed photos (except iNs, as mentioned above). The figures in the article have been modified to enhance clarity, ensuring that all representative pictures are processed similarly.

### 7.9. Mitochondrial Mass

To quantify the mitochondrial mass, the mitochondria were stained using the mitochondrial dye MitoTracker™ Green FM. MitoTracker™ Green FM does not rely on MMP for its fluorescence signal, consequently allowing for a correlation between the fluorescent signal of MitoTracker™ Green FM and the mitochondrial mass. The CellTracker™ Blue CMAC Dye was used to standardize the fluorescence signal to identify the cellular surface area. All cell models were stained with MitoTracker™ Green FM (0.1 μM) and CellTracker™ Blue CMAC Dye (5 μM) for 1 hour at 37 °C and 5 % CO_2_. Subsequently, the cells underwent two washes with HBSS. The fluorescence signals of MitoTracker™ Green FM at 490 nm (excitation) and 516 nm (emission), as well as CellTracker™ Blue CMAC Dye at 353 nm (excitation) and 466 nm (emission), were measured at the same time using the Cytation 3 Cell Imaging Multi-Mode Reader. The mitochondrial mass was determined by calculating the mitochondria-to-cellular mass ratio.

### 7.10. Telomere length

#### 7.10.1. DNA isolation

The DNA was isolated according to the FlexiGene DNA Handbook using the FlexiGene^®^ DNA KIT (250). Briefly, a quantity of 1-2 x 10^6^ cells was combined with 750 μl FG1 buffer and centrifuged at 10000 x g for 20 seconds. In the next step, 150 μl FG2/GIAGEN protease was mixed into the tube and then incubated at 65°C for 5 min. Afterward, 150 μl of isopropanol (100 %) was added and vortexed until a DNA precipitate was observed. Centrifugation followed this step for 5 min at 10000 *x* g. The supernatant was tossed, and the pellet was resuspended in 150 μl of 70 % ethanol by vortexing for 5 sec. After discarding the supernatant, the DNA pellet was air-dried until the liquid evaporated. The DNA pellet was then dissolved in 200 μl buffer FG3 by vortexing for 5 sec at low speed, followed by incubation for 60 min at 65 °C in a heating block. Finally, the DNA content was assessed using the Take3^TM^ Microvolume Plate in combination with the Cytation 3 Cell Imaging Multi-Mode Reader, and the DNA samples were stored at -20°C until further use.

#### 7.10.2. Telomere length determination by qPCR

The telomere length was measured by quantitative polymerase chain reaction (qPCR) according to approaches described previously [154–157]. To determine the telomere length, the T/S ratio (telomere repeat copy number (TELO) to single-copy gene number (SCG), which represents a relative measure of the telomere length, was accessed. The TELO and SCG were detected for each DNA sample. Table 4 lists the forward and reverse primers of TELO and SCG (ß-globin). A master mix was prepared for the qPCR. The composition of one reaction of TELO master mix was 1.08 ul teloF primer (54 nM), 1,6 μl teloR primer (80 nM), 5,32 μl H2O, 10 μl of GoTaq^®^ qPCR Master Mix, and 0.2 μl Supplemental CXR Reference Dye and for SCG master mix was 1,6 μl ß-globinF primer (80 nM), 1,6 μl ß-globinR primer (80 nM), 4,8 μl H2O 10 μl of GoTaq^®^ qPCR Master Mix, and 0.2 μl Supplemental CXR Reference Dye. The master mixes of TELO or SCG (18 μl per well) were loaded into 96-well PCR plates. Afterwards, 2 μl of the DNA sample was added to the corresponding wells at a final concentration of 10 ng/μl. The qPCR settings were first the initial denaturation at 95°C for 10 min, followed by cycling of 50 repeats containing 1.) 10-sec hold at 95°C and 2.) 60-sec hold at 58 °C. The Ct values were exported with the SABiosciences PCR Array Data Analysis Software and analyzed with the comparative Ct method (2^-ΔΔCt^) relative to the internal control to be represented as the T/S ratio. As internal control, a mixture of the HFs DNA samples was used.

**Table 4:**
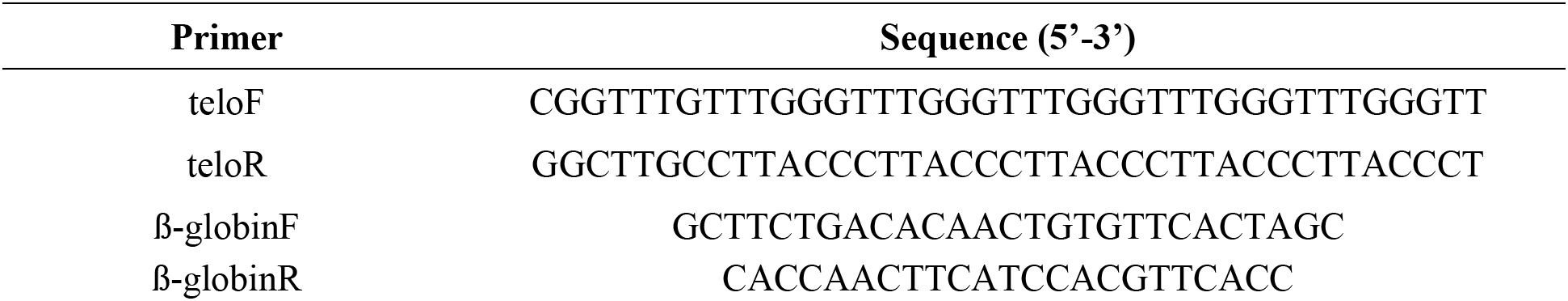
Forward and reverse primer sequences of TELO and SCG.

### 7.11. Telomerase activity

Telomerase Repeat Amplification Protocol (TRAP) was performed to determine the telomerase activity [158–160]. This TRAP assay is based on three steps: extension, amplification, and detection. The extension step is characterized by adding telomeric repeats to the DNA by the telomerase activity during a defined time frame. In the amplification and detection part, the extension is amplified and detected by qPCR. Therefore, a defined amount of cells (HFs and iPSCs: 10 000 cells; iNs and iPSCsN: 100 000 cells) were lysed in ice-cold NP-40 lysis buffer (10 mM Tris-HCl (pH 8.0); 1 mM MgCl_2_(6 H_2_O); 1 mM EDTA, 1 % (v/v) NP-40, 0.25 mM sodium deoxycholate, 10 % (v/v) glycerol, 150 mM NaCl, 5 mM 2-mercaptoethanol, and 0.1 mM AEBSF) and incubated for 30 min on ice. Following the incubation, the samples were centrifuged for 20 min at 14’000 rpm at 4°C. Next, the supernatant was collected and further utilized in the TRAP assay. In this case, the sample was snap-frozen and then stored at -80°C. A master mix containing 0.2 μl ACX-primer (300 nM), 0,2 μl TS-Primer (300 nM), 7,6 μl DNase/RNase free water 10 μl of GoTaq® qPCR Master Mix, and 0.2 μl Supplemental CXR Reference Dye per one reaction was prepared. The primer sequences are listed in Table 5. The master mix (18 μl per well) was loaded in 96-well PCR plate. Followed by 2 μl of the lysate was added to the corresponding well. A positive control of HEK293 cells in the range of 10’000 cells to 640’000 cells was measured on every PCR plate. The preparation of the TRAP assay was done on ice. The real-time PCR was performed using the Step One Plus system. The settings of the measurement were for the extension at 25°C for 40 min, next to the deactivation of telomerase at 95°C for 10 min, followed by a cycling of 40 repeats containing 1.) 30-sec hold at 95°C, 2.) 30-sec hold at 52 °C and 3.) 45-sec at 72°C. The Ct values were exported using the SABiosciences PCR Array Data Analysis Software. The final data were represented as the normalization to the positive control.

**Table 5:**
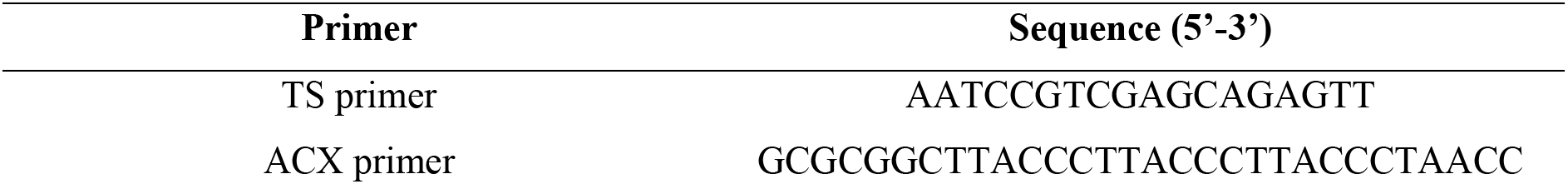
Primer sequences of TRAP assay.

### 7.12. Gene expression

#### 7.12.1. RNA isolation

The RNA was isolated according to the manufacturing protocol of the RNeasy Mini Kit. The cells were harvested in 350 μl RLT lysis buffer on ice, and then 350 μl of 70 % ethanol was added. The solution was transferred to the RNeasy spin column with a collecting tube and centrifuged at ≥ 10’000 rpm for 15 sec. The flow-through was discarded. Next, the column was washed with 700 μl Buffer RW1 and followed twice with 500 μl RPE. Then, the column was tried at 14000g for 1 min. Finally, the RNA was collected with 30 μl RNase-free water by waiting 1 min and then centrifuging for 15 sec at ≥ 8000g to collect the RNA. The centrifuge was set up at 4°C during the RNA isolation process. The RNA content was assessed by using a Nanodrop 1000 spectrophotometer.

#### 7.12.2. Reverse Transcription and Gene expression

The extracted RNA was transformed to cDNA using the RT^2^ First Strand Kit [161] by following the manufactural instructions to measure the gene expression in the neuronal cells. Next, the gene expression was detected using the Custom RT^2^ Profiler PCR Arrays with the RT^2^ SYBR Green ROX qPCR Master Mix according to the manufacturer’s guidelines. The RT^2^ Profiler PCR Arrays were pre-coated with custom-chosen primers (primers listed in Table 6). Step One Plus system was used to perform the quantitative real-time PCR. The CT values (automatically generated by the Step One Plus software) were exported and analyzed with the GeneGlobe Data Analysis Centre Software and MetaboAnalyst 5.0.

**Table 6:** List of genes assessed with the Custom RT^2^ Profiler PCR Arrays (Qiagen). AKT1: Serine/threonine-protein kinases 1; BCL2: B-cell lymphoma 2; CAT: Catalase; DNM1L: Dynamin-related protein 1 (Drp1); FIS1: Mitochondrial Fission Protein 1;FOXO1: Forkhead box protein O1; GPX1: Glutathione peroxidase 1; LDHA: Lactate dehydrogenase A (LDHA); MFN1: Mitofusin-1; MFN2: Mitofusin-2; MTOR: Mammalian target of rapamycin; NFE2L2: Nuclear factor erythroid-derived 2-like 2 (Nrf2); OPA1:Optic atrophy 1; PARP1:Poly (ADP-ribose) polymerase 1; PFKFB3: 6-phosphofructo-2-kinase/fructose-2,6-biphosphatase 3; PPARGC1A: Peroxisome proliferator-activated receptor gamma coactivator 1-alpha (PGC-1α); PRKAA2: AMP-activated protein kinase (AMPK); SIRT1: Silent mating type information regulation 2 homolog 1; SIRT3: Silent mating type information regulation 2 homolog 3; SOD1:Superoxide dismutase 1;SQSTM1: Sequestosome 1 (p62); TFAM: Mitochondrial transcription factor; TP53: Transformation-related protein 53 (p53); UCP2: Mitochondrial uncoupling protein 2

| Gene symbol | Assay Nr. | GeneBank |
| --- | --- | --- |
| PPARGC1A | PPH00461F | NM_013261 |
| TFAM | PPH09934B | NM_003201 |
| SIRT3 | PPH22989A | NM_012239 |
| PARP1 | PPH00686B | NM_001618 |
| PFKFB3 | PPH00024F | NM_004566 |
| TP53 | PPH00213F | NM_000546 |
| FIS1 | PPH19947A | NM_016068 |
| OPA1 | PPH12084A | NM_130837 |
| MFN2 | PPH18422A | NM_014874 |
| MFN1 | PPH20215A | NM_033540 |
| DNM1L | PPH13763A | NM_005690 |
| BCL2 | PPH00079B | NM_000633 |
| SOD1 | PPH00234B | NM_000454 |
| CAT | PPH00420B | NM_001752 |
| GPX1 | PPH00154F | NM_000581 |
| MTOR | PPH02311D | NM_004958 |
| AKT1 | PPH00088B | NM_005163 |
| SIRT1 | PPH02188A | NM_012238 |
| UCP2 | PPH00476F | NM_003355 |
| NFE2L2 | PPH06070A | NM_006164 |
| PRKAA2 | PPH15207A | NM_006252 |
| LDHA | PPH02047H | NM_005566 |
| SQSTM1 | PPH02107A | NM_003900 |
| FOXO1 | PPH01964F | NM_002015 |
| GAPDH | PPH00150F | NM_002046 |

### 7.13. Normalization

All data were normalized to the cell number determined by DAPI staining, except for the NAD^+^/NADH, TA, TL, and gene expression. After each experiment, the cells were fixed with 4 % formaldehyde for 10 minutes and washed twice with PBS. Followed by staining with DAPI for 5 min; the cells were washed once with PBS. The cell number was determined using the cell analyzing settings on the Gen5 Image 3.11 software of the Citation 3 Cell Imaging Multi-Mode Reader. The average cell count from a parallel running plate was utilized for the cellular ATP level. The NAD^+^ and NADH values were normalized to the protein level. The mitochondrial mass was standardized by dividing to the cell surface.

### 7.14. Statistical Readout

Data management was performed using Graph Pad Prism 9 software, Excel software, R-software, and metaboanalyst.ca website. The presented data indicate the mean ± SEM and normalized parato the representative young cell types. The statistical significance analysis was done to compare two groups by student unpaired t-test with the p-values: * p < 0.05, ** p < 0.01, and *** p < 0.001 by comparing young versus aged. PCA was conducted to visually represent the primary directions that most accurately characterize the variability in our dataset without explicitly referring to group labels. A heat map was generated from a hierarchical cluster analysis using Euclidean distance and Ward’s linkage method. To generate the radar charts for the donor comparison, the statistical software R (version 4.2.1), including the packages fmsb (version 0.7.5) [162], was used.

## Supporting information

Expanded View

## 1. Abbreviation

^-^OH: Hydroxyl radical
°O_2−_: Superoxide anion radicals
AKT1: Serine/threonine-protein kinases 1
AMPK: AMP-activated protein kinase
AR: Aspect ratio
ATP: Adenosine triphosphate
AW: Area weigthed form factor
BCL2: B-cell lymphoma 2
CAT: Catalase
DHR: Dihydrorhodamine 123
DNM1L: Dynamin-related protein 1 (gene)
DRP1: Dynamin-related protein 1 (protein
ECAR: Extracellalur acidification rate
em: Emission
ex: Excitation
F: Female
FF: Form factor
FIS1: Mitochondrial fission protein 1
FOXO1: Forkhead box protein O1
Gly.: Glycolytic
GM: Growing medium
GPX1: Glutathione peroxidase 1
H^+^: Protons
H_2_O_2_: Hydrogen peroxide
HEK293: Human embryonic kidney 293
HFs: Human fibroblasts
iNs: Directly converted neurons
iNs: Induced neurons
iPSCs: Induced pluripotent stem cells
iPSCsNs: Induced pluripotent stem cells derived neurons
LDHA: Lactate dehydrogenase A
LMNA: Lamin A
Max.: Maximal
MFN1: Mitofusin-1
MFN2: Mitofusin-2
min: Minute
mito.: Mitochondria mitogenesis Mitochondrial biogenesis<colcnt=2>
MMP: Mitochondrial membrane potential
Morph.: Morphology
mTOR: Mammalian target of rapamycin
NAD: Nicotinamide adenine dinucleotide
NEM: Neural expansion medium
NFE2L2: Nuclear factor erythroid-derived 2-like 2 (Nrf2)
NIM: Neural induction medium
NK: Neuronal conversion
Norm.: Normalised
NPCs: Neural progenitor cells
NRF2: Nuclear factor erythroid-derived 2-like 2
OCR: Oxygen consumption rate
ONOO^-^: Peroxynitrite
OPA1: Optic atrophy 1
OXPHOS: Oxidative phosphorylation
p53: Transformation-related protein 53
p62: Sequestosome 1
PARP1: Poly (ADP-ribose) polymerase 1
PCA: Principal component analysis
PFKFB3: 6-phosphofructo-2-kinase/fructose-2,6-biphosphatase 3
PGC-1α: Peroxisome proliferator -activated receptor gamma coactivator 1 alpha
PPARGC1A: Peroxisome proliferator-activated receptor gamma coactivator 1-alpha (PGC-1α)
PRKAA2: AMP-activated protein kinase
qPCR: Quantitative PCR
Resp.: Respiration
ROS: Reactive oxygen species
SCG: Single-copy gene number
SD: Standard deviation
SIRT1: Silent mating type information regulation 2 homolog 1
SIRT3: Silent mating type information regulation 2 homolog 3
SOD1: Superoxide dismutase 1
SQSTM1: Sequestosome 1
TA: Telomerase activity
TCA: Tricarboxylic acid
TELO: Telomere repeat copy number
TFAM: Mitochondrial transcription factor A
TL: Telomere length
TMRM: Tetramethyl rhodamine, methyl ester, and perchlorate
TP53: Transformation-related protein 53
TRAP: Telomerase repeat amplification protocol
TRAP: Telomerase repeated amplification protocol
UCP2: Mitochondrial uncoupling protein 2
UNA: plvx-ubc-rtta-Ngn2:2A:ASCL1
Y-27632: Rock-inhibitor

## 8. Acknowledgements

We are grateful thank Stella Stefanova, Svitlana Malysheva and Janine Bögli of the FACS core facility at the Biozentrum in Basel for the great support and measurements. Further, we thank Fides Meier for general laboratory coordination. This work was supported by the Swiss National Science Foundation 31003A-179294 (A.E.), the Novartis Foundation for Medical Research 18C143 (A.E.), Innovative Medicines Initiative Joint Undertaking 115439 (Z.C.), European Union’s Seventh Framework Programme FP7/2007-2013 (Z.C.), and EFPIA companies (Z.C.).

## 10. Author contributions

**Nimmy Varghese:** Formal analysis; investigation; methodology; visualization; writing – original draft. **Leonora Szabo:** Methodology; investigation. **Zameel Cader:** Resources; methodology**. Imane Lejri:** Investigation. **Amandine Grimm:** Methodology; supervision. **Anne Eckert:** Conceptualization; Resources; supervision; funding acquisition; project administration.

## 11. Data Availability Section

The data that support the findings of this study will be openly available after the peer-review process.

## 12. Expanded view

Please refer to the attached file.

