## Supplementary material for "Preservation of an Aging-Associated Mitochondrial Signature in Advanced Human Neuronal Models": Expanded View

**Expanded View Table 1: Determination of $NAD^+$ to NADH ratio.** Data unit: $NAD^+$ [nM] or NADH [nM] /norm. to protein concentration [mg/ml]. Abbreviation: HFs: Human fibroblasts; iNs: directly induced neurons; iPSCsNs: induced pluripotent stem cells (iPSCs) derived neurons

| | **HFs** | | **iNs** | | **iPSCsNs** | |
|---|---|---|---|---|---|---|
| | **Young** | **Aged** | **Young** | **Aged** | **Young** | **Aged** |
| **$NAD^+$ (mean)** | 218.85 | 41.85 | 12.51 | 7.87 | 112.39 | 75.71 |
| **NADH (mean)** | 30.82 | 75.21 | 23.57 | 34.24 | 29.02 | 60.64 |
| **Ratio** | 7.10 | 0.56 | 0.53 | 0.23 | 3.87 | 1.25 |

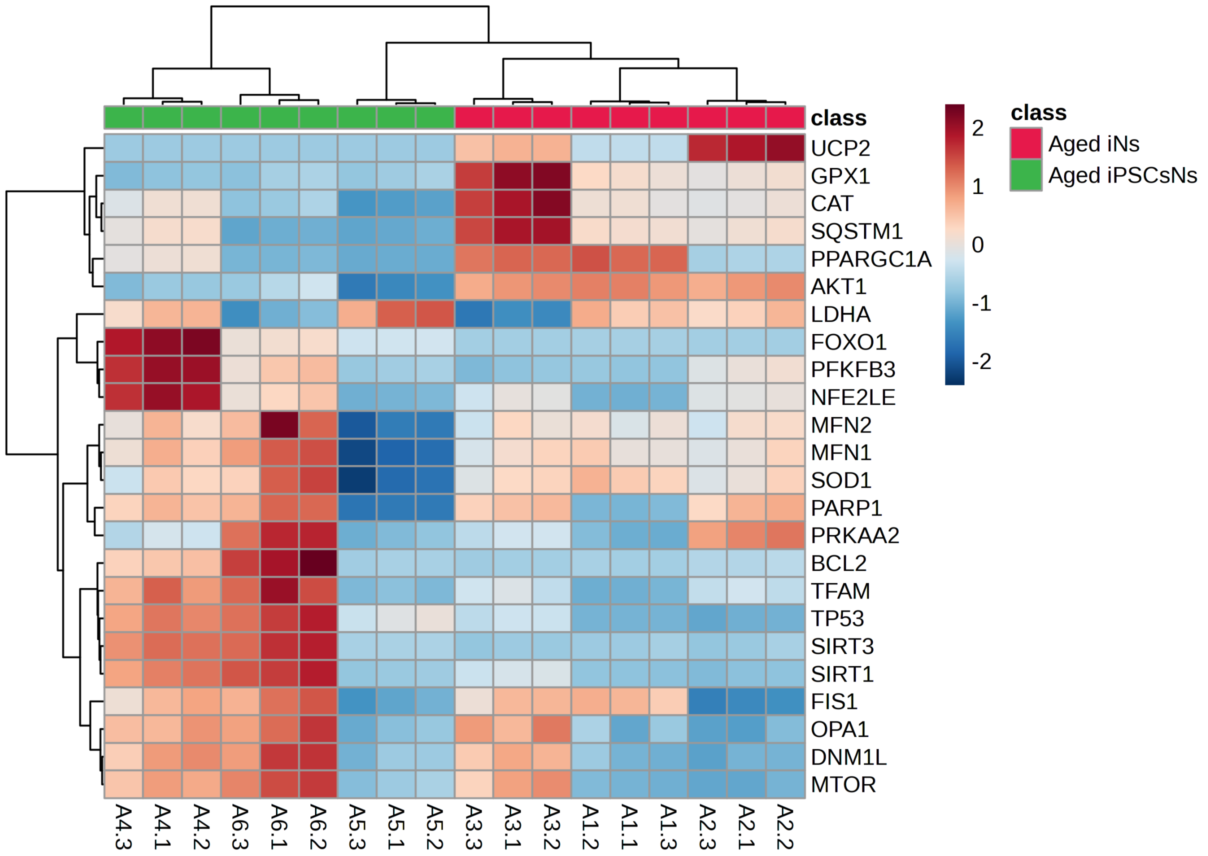


**Expanded Figure 1: Investigation of differences in aged iNs versus aged iPSCsNs using quantitative real-time PCR analysis.** Clustering results are shown as a heatmap of aged iNs vs. aged iPSCsN (distance measured using Euclidean, and clustering algorithm using ward.D).

**Data information:** The figure was generated on metaboanalyst.ca. The represented values were normalized by autoscaling (mean-centered and divided by the standard deviation of each variable). The data were represented as Gene expression ($2^{(\text{-Avg.(Delta(Ct))})}$) by using the Houskeeping gene GAPDH. Abbreviation: AKT1: Serine/threonine-protein kinases 1; BCL2: B-cell lymphoma 2;CAT: Catalase; DNM1L: Dynamin-related protein 1 (DRP1); FIS1: Mitochondrial Fission Protein 1;FOXO1: Forkhead box protein O1; GPX1: Glutathione peroxidase 1; iNs: Induced neurons; iPSCs: induced pluripotent stem cells; iPSCsNs: induced pluripotent stem cells (iPSCs) derived neurons; LDHA: Lactate dehydrogenase A (LDHA); MFN1: Mitofusin-1; MFN2: Mitofusin-2; MTOR: Mammalian target of rapamycin; NFE2L2: Nuclear factor erythroid-derived 2-like 2 (NRF2); OPA1: Optic atrophy 1; PARP1: Poly (ADP-ribose) polymerase 1; PFKFB3: 6-phosphofructo-2-kinase/fructose-2,6-biphosphatase 3; PPARGC1A: Peroxisome proliferator-activated receptor gamma coactivator 1-alpha (PGC-1α); PRKAA2: AMP-activated protein kinase (AMPK); SIRT1: Silent mating type information regulation 2 homolog 1; SIRT3: Silent mating type information regulation 2 homolog 3; SOD1: Superoxide dismutase 1;SQSTM1: Sequestosome 1 (p62); TFAM: Mitochondrial transcription factor; TP53: Transformation-related protein 53 (p53); UCP2: Mitochondrial uncoupling protein 2. Source data will be available online for all figures. Please refer to the data availability section.

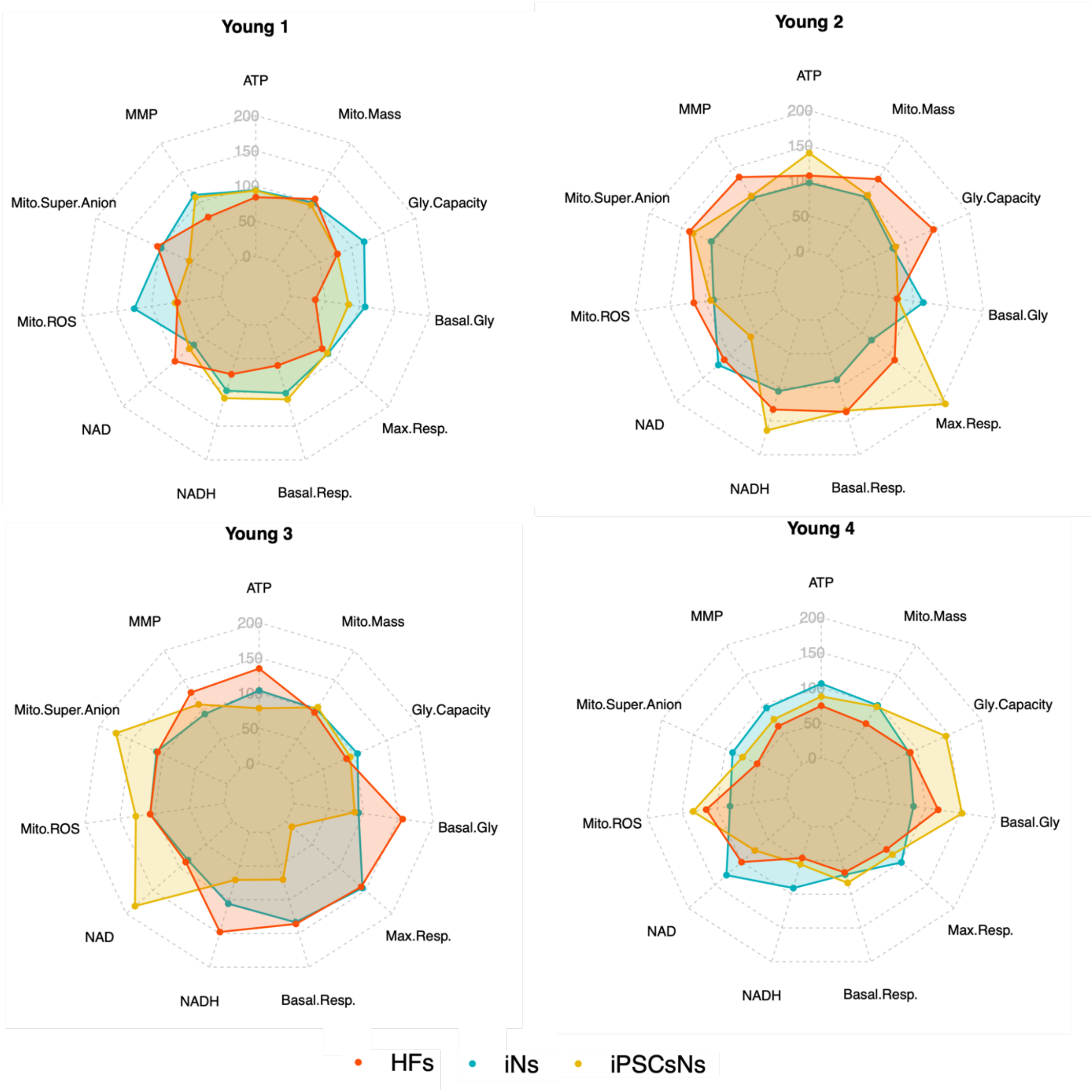


**Expanded Figure 2: The radar plot summarizes values from each young donor and shows the different cell models of the same donor in a single graph.** The radar plot was generated with the R-software. The data used to generate the radar plot are represented as % a percentage of the mean young and are listed in the EV Table 2. Abbreviation: Gly: Glycolytic; HFs: Human fibroblasts; iNs: directly induced neurons; iPSCsN: induced pluripotent stem cells (iPSCs) derived neurons; Max: maximal; Mito: Mitochondria, MMP: mitochondrial membrane potential; Resp: Respiration. Source data will be available online for all figures. Please refer to the data availability section.

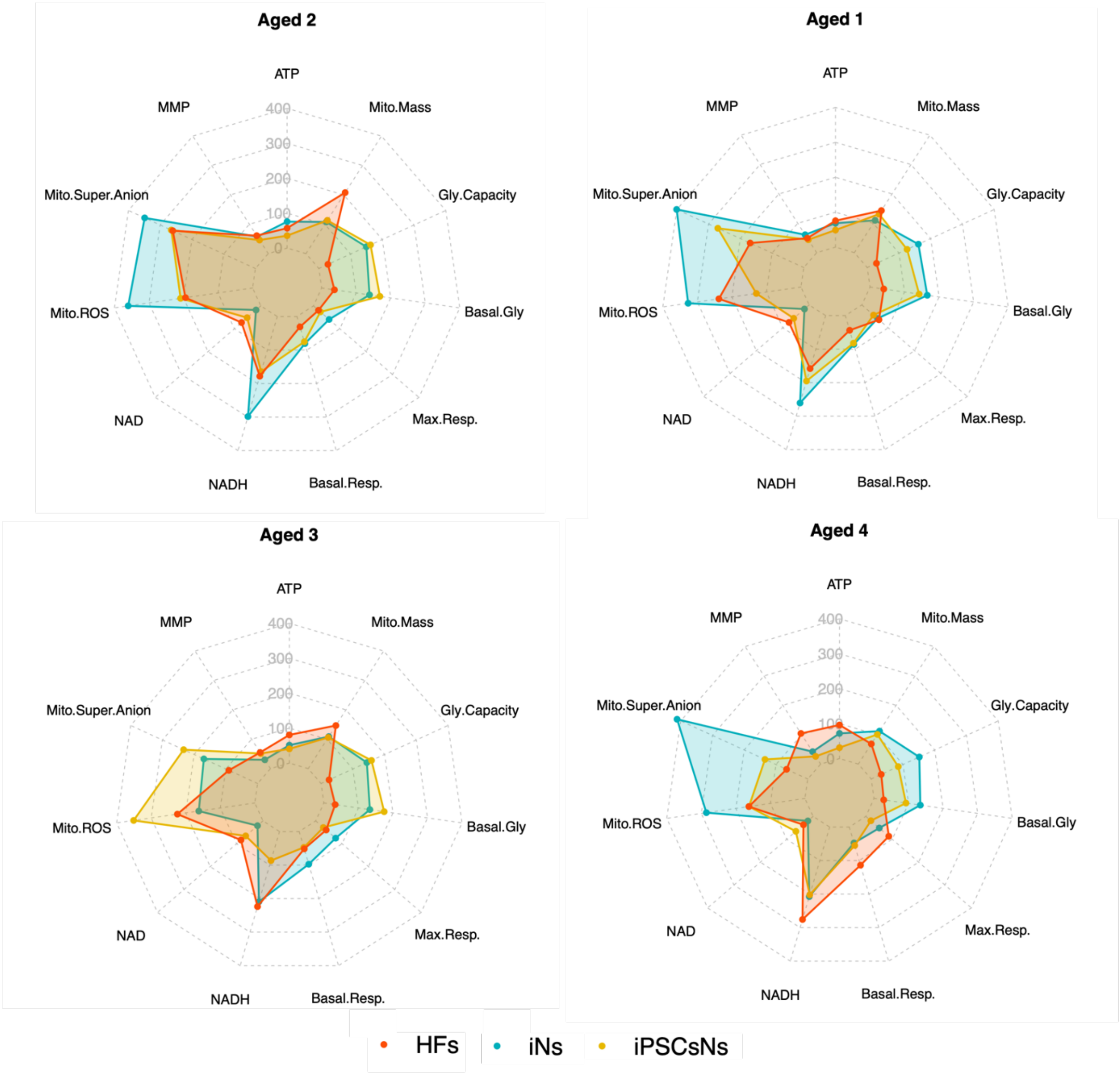


**Expanded Figure 3: The radar plot summarizes values from each aged donor and represents the different cell models in the same graph.** The radar plot or spider web were generated with the R-software. The data used to generate the radar plot are the represented as % percentage to the mean young and are listed at the (EV Table 2). Abbreviation: Gly: Glycolytic; HFs: Human fibroblasts; iNs: directly induced neurons; iPSCsN: induced pluripotent stem cells (iPSCs) derived neurons; Max: maximal; Mito: Mitochondria, MMP: mitochondrial membrane potential; Resp: Respiration. Source data will be available online for all figures. Please refer to the data avaibility section.

**Expanded View Table 2: The data used to construct the radar plot for donor comparison.** The data depict normalized values in relation to the corresponding young cells (100 %). Abbreviation: Gly: glycolytic; HFs: Human fibroblasts; iNs: directly induced neurons; iPSCsNs: induced pluripotent stem cells (iPSCs) derived neurons; Max: maximal; Mito: mitochondria, MMP: mitochondrial membrane potential; Resp: respiration.

| | Group | ATP | MMP | Mito. ROS | Mito. Superoxide Anion | $NAD^+$ | NADH | Basal. Resp. | Max.Resp. | Basal.Gly. | Gly. Capacity. | Mito.Mass |
|---|---|---|---|---|---|---|---|---|---|---|---|---|
| HFs | Young1 | 93.63 | 112.88 | 97.85 | 124.60 | 66.73 | 97.46 | 101.21 | 85.62 | 107.04 | 119.36 | 99.81 |
| | Young2 | 97.04 | 99.29 | 103.11 | 87.32 | 121.04 | 105.77 | 88.79 | 67.04 | 113.63 | 80.29 | 100.69 |
| | Young3 | 103.60 | 92.81 | 110.43 | 106.46 | 83.87 | 105.89 | 133.47 | 144.31 | 92.77 | 103.70 | 101.21 |
| | Young4 | 105.73 | 94.00 | 88.61 | 81.09 | 128.36 | 90.88 | 70.97 | 100.70 | 82.54 | 87.45 | 98.25 |
| | Aged1 | 68.49 | 61.10 | 400.63 | 326.74 | 17.83 | 260.76 | 86.03 | 59.29 | 165.80 | 161.13 | 110.08 |
| | Aged2 | 75.83 | 57.94 | 348.91 | 360.27 | 17.94 | 298.40 | 80.74 | 59.27 | 138.58 | 149.59 | 107.80 |
| | Aged3 | 51.24 | 30.06 | 169.35 | 162.23 | 20.87 | 209.34 | 97.40 | 75.43 | 133.05 | 144.13 | 108.92 |
| | Aged4 | 72.02 | 42.31 | 411.57 | 285.30 | 19.85 | 207.46 | 47.60 | 50.91 | 133.99 | 151.44 | 112.20 |
| iNs | Young1 | 93.07 | 108.99 | 53.94 | 65.82 | 74.86 | 108.66 | 110.47 | 84.55 | 83.33 | 77.75 | 95.71 |
| | Young2 | 139.34 | 102.37 | 131.74 | 91.62 | 60.11 | 163.81 | 134.51 | 205.80 | 77.28 | 85.07 | 103.72 |
| | Young3 | 78.30 | 109.00 | 173.57 | 127.10 | 183.53 | 70.71 | 70.15 | 11.48 | 87.16 | 92.39 | 104.25 |
| | Young4 | 87.48 | 74.55 | 72.95 | 134.04 | 74.77 | 55.59 | 83.28 | 83.73 | 152.23 | 144.79 | 96.06 |
| | Aged1 | 48.91 | 44.10 | 270.76 | 129.21 | 58.55 | 195.27 | 82.42 | 44.29 | 142.24 | 125.55 | 131.27 |
| | Aged2 | 35.58 | 46.19 | 265.05 | 208.46 | 51.73 | 164.61 | 75.38 | 26.39 | 168.64 | 162.84 | 113.16 |
| | Aged3 | 40.69 | 51.50 | 233.05 | 350.78 | 65.62 | 86.64 | 47.32 | 29.01 | 174.53 | 158.82 | 105.98 |
| | Aged4 | 31.29 | 26.29 | 134.74 | 162.05 | 65.42 | 202.27 | 55.07 | 19.05 | 92.46 | 85.21 | 100.91 |
| iPSCsNs | Young1 | 83.74 | 75.30 | 103.65 | 62.45 | 102.05 | 73.17 | 60.12 | 74.98 | 35.59 | 77.59 | 105.95 |
| | Young2 | 107.26 | 134.42 | 137.32 | 115.84 | 109.93 | 132.75 | 136.31 | 110.51 | 75.82 | 143.94 | 130.97 |
| | Young3 | 134.79 | 129.19 | 108.73 | 106.53 | 88.03 | 147.82 | 135.83 | 141.86 | 156.02 | 86.46 | 95.63 |
| | Young4 | 74.21 | 63.04 | 50.30 | 115.18 | 99.99 | 46.26 | 67.73 | 72.65 | 117.92 | 89.19 | 67.44 |
| | Aged1 | 75.66 | 49.40 | 168.89 | 237.74 | 76.45 | 158.16 | 43.94 | 64.78 | 39.78 | 29.14 | 143.30 |
| | Aged2 | 57.00 | 61.18 | 260.39 | 194.42 | 72.91 | 178.10 | 30.92 | 19.11 | 37.03 | 28.65 | 207.65 |
| | Aged3 | 80.62 | 54.93 | 90.65 | 224.14 | 83.48 | 223.79 | 51.63 | 39.69 | 32.67 | 25.12 | 146.56 |
| | Aged4 | 96.15 | 104.31 | 66.69 | 162.18 | 36.62 | 275.72 | 113.85 | 86.99 | 28.13 | 31.50 | 68.72 |
